# Investigating the neural effects of typicality and predictability for face and object stimuli

**DOI:** 10.1101/2023.10.24.563242

**Authors:** Linda Ficco, Chenglin Li, Jürgen M. Kaufmann, Stefan R. Schweinberger, Gyula Z. Kovács

## Abstract

The brain calibrates itself based on the past stimulus diet, which makes frequently observed stimuli appear as typical (as opposed to uncommon stimuli, which appear as distinctive). Based on predictive processing theory, the brain should be more “prepared” for typical exemplars, because these contain information that has been encountered frequently, and allow to economically represent items of that category. Thus, one could ask whether predictability and typicality of visual stimuli interact, or rather act in an additive manner. We adapted the design by Egner and colleagues (2010), who used cues to induce expectations about stimulus category (face vs. chair) occurrence during an orthogonal inversion detection task. We measured BOLD responses with fMRI in 35 participants. First, distinctive stimuli always elicited stronger responses than typical ones in all ROIs, and our whole-brain directional contrasts for the effects of typicality and distinctiveness converge with previous findings. Second and importantly, we could not replicate the interaction between category and predictability reported by Egner et al. (2010), which casts doubt on whether cueing designs are ideal to elicit reliable predictability effects. Third, likely as a consequence of the lack of predictability effects, we found no interaction between predictability and typicality in any of the four tested regions (bilateral fusiform face areas, lateral occipital complexes) when considering both categories, nor in the whole brain. We discuss the issue of replicability in neuroscience and sketch an agenda for how future studies might address the same question.

## 1. Introduction

### 1.1 Perceptual Spaces in Visual Perception

If one asks you to imagine the picture of a chair, chances are that it would be made of wood, have four legs, be rather symmetrical, have a simple backrest and maybe a pillow. Within any visual category, some exemplars are more representative than others, and are perceived as “typical” (1). The perception of typicality depends on the fact that exemplars are not represented independently in our mind (and brain). Any chair exemplar is remembered and processed in relation to other similar-looking items, seen in similar contexts, with the same function, size, material, and so on. Perceptual categories, such as “chairs”, can be thought of as low-dimensional perceptual spaces of various exemplars (2–4), with minimal differences and maximal similarities across exemplars ((1), p. 491).

In such models, typical exemplars share most features with each other (5), have more average features (6), stand out of the category less (7), and summarize its dimensions best (5,8). Importantly, the perception of typicality is flexible: it is influenced by the current perceptual diet of the observer (9) and its stability depends on the duration and amount of previous exposure (see (1) for a discussion).

Faces are thought to be represented in such space (6,10,11). We are exposed to thousands of faces throughout our life (12), we develop expertise for this category (13,14). Our face diet shapes the prototype in reference to which new faces are encoded constantly (15–18), and people with allegedly comparable face spaces tend to form coherent typicality judgements about the same faces (19).

A long-standing debate exists regarding different face-space models (11,20,21). In exemplar-based models faces are encoded based on their similarity, at relative distances from each other (6,22). Conversely, in norm-based models, faces are encoded with respect to a weighted average of all seen faces (23). Notably, norm-based coding appears to extend beyond facial stimuli to non-face objects as well (24–27). When comparing the two models explicitly, the norm-based model seems to account for a number of findings – e.g., responses in the fusiform face area – better than the exemplar-based model (28). However, note that other works comparing these two models, and using visual stimuli for which participants have acquired expertise (e.g., faces, real objects and abstract shapes), seem to find support for both (21,24,29,30).

### 1.2 Behavioural and Neural Effects of Typicality

Despite different existing operationalizations of typicality (e.g., different kinds of ratings, via morphing, etc…), its effects are relatively uncontroversial: Neural and behavioural data generally show that typical stimuli are easier to *process* (that is, performing mental operations, such as detection or classification, on them) but harder to *encode* (i.e., store in a mental representation). Behaviourally, typical stimuli are categorized faster and more accurately than distinctive ones (24,25,31–34). Moreover, typical stimuli are detected faster than distinctive ones (35), as suggested by detection advantages for own-“race” (typical), as compared to other-“race” faces (6,36). However, typicality hampers the recognition of specific exemplars: distinctive faces are recognized more accurately (37,38,47,39–46) and faster (34,37–39,46) but see (48) than typical faces. This is possibly due to poorer encoding of typical stimuli (49), which are more similar to each other and can be confused easier with each other, leading to false positive answers by worse pattern separation processes (50).

Several neuroimaging techniques have been used to map perceptual spaces (reviewed in (51). Neuroimaging findings suggest that typical stimuli require i) smaller brain metabolic changes and ii) reduced configural processing, as compared to distinctive stimuli. Specifically, fMRI studies report increased brain responses to distinctive stimuli, both in the visual (28,52,53) and in the auditory modality (54,55). Conversely, electrophysiological studies tend to use event-related potentials (ERPs). In ERP studies, face typicality consistently affects the amplitude of the occipito-temporal P200 component, with larger amplitudes for typical, as compared to distinctive stimuli (10,56–58). Object typicality potentially affects occipito-temporal ERPs even at latencies < 200 ms (59); studies using objects or animal images report larger P200 in frontal regions to distinctive as compared to typical objects (31) and shorter latencies of the P300 component for typical stimuli (60). These findings parallel reaction time data (60) and could indicate that typical stimuli require less attention and lower feature encoding (61).

Importantly, and although it should be kept in mind that the precise definition of typicality may vary across individual studies, neural and behavioural typicality effects appear to be related, and their relationship cannot be explained by the mere physical similarity of stimuli (52).

### 1.3 Typicality Effects under the Predictive Coding Framework: The Current Study

Visual perception does not only capitalize on dimensionality reduction (as discussed above), but also on the prediction of upcoming sensory inputs (62). According to the theory of predictive processing (63–65), the brain attempts to predict the input received from lower processing levels, by generating an internal model of the input at each level of stimulus processing (be it single neurons, neural populations, or network hubs) (66,67). The degree to which this internal model differs from the incoming input represents the prediction error (i.e., the mismatch between actual and predicted input; (68)), which is then transferred to higher levels (67). These recursive model-error comparison loops tend to minimize prediction error (69), and ultimately allow the individual to bear the most accurate and up-to-date models of the sensory world. In other words, perception is not a passive process, but an active attempt of the brain to guess the latent causes of a sensory input best, informed by prior knowledge (70,71).

Visual prototypes seem to be just this: representations “summarizing” stimuli from a particular category best, based on those stimuli we frequently experienced in the past. Potentially, this theory elegantly explains the neural and behavioural effects of stimulus typicality as well: typical stimuli elicit less prediction error than distinctive ones when compared to a prototype, thus leading to faster detection and categorization, and lower neural responses in regions processing stimulus configuration and category. However, to date systematic research on the combined effects of predictability and typicality is missing. To address this important theoretical and empirical gap, one should test to which degree typicality and predictability *interact*, or act in an *additive manner.* An interaction between predictability and typicality would imply a common neural mechanism underlying both effects, whereas an additive effect would be more consistent with separable mechanisms to compute stimulus probability and typicality (a logic similar as that used in (72,73). Accordingly, if two processes are independent, the expression of one effect should not change for different levels of the other (i.e., the two effects should just add up). Instead, if there is an additional increase in one effect for one level of the other that is not explained by a simple addition of the two effects, one can infer an (at least partial) dependency of the mechanisms underlying the two effects.

To test this, we presented face or chair stimuli, preceded by a cue which signalled the participants the probability of the occurrence of a stimulus category (“predictability” here thus indicates expected temporal association between a stimulus category and a cue preceding the stimulus). Importantly, orthogonal to this predictability modulation, we also manipulated stimulus typicality by presenting typical and distinctive stimulus exemplars (so, typicality here refers to the distance of each stimulus from an average exemplar of the same category – see Section 2.2). We replicated a design that produced an interaction between category and predictability in a previous study (74), and that could allow us to elegantly test the presence of an interaction. Another important goal of the present work is to assess the replicability of this effect, especially since a recent, large study (75) showed it is challenging to produce neural predictability effects with cueing, despite effective behavioural training.

## 2. Materials and Methods

### 2.1 Participants

We recruited thirty-five adult participants (22 females, one diverse; mean age = 24.4 years; SD = 4.0 years; two left-handed, one ambidextrous), with normal or corrected-to-normal vision and with no reported neurological condition (one participant reported a diagnosis of Asperger Syndrome, their data were retained in the analysis). The measurements were performed between May and August 2022. The main experimenter had access to identifying information from participants and took care of making their data anonymous for the analyses. We ensured that each had > 10 years-long exposure to Caucasian/European/White faces. The sample size was determined by a power analysis with the R package *Superpower* (76). We calculated the number of participants necessary to detect a medium effect size (f = 0.25) for the interaction of cue-induced predictability and category, to achieve a power of 0.80 at the standard .05 alpha error probability. This was calculated based on the mean beta estimates in different conditions reported in Figure 3 of (74). We chose this effect for our power analysis because we had no expectations about the size of the interaction of interest, so we preferred to calibrate our sample size to target a known effect. This led to a sample size that nearly doubled the sample reported in (74), who recruited 17 participants. Participants could choose between monetary compensation and a small 3D model of their own brain for their participation. Before the experiment, all participants received information about the experimental procedures and provided their written informed consent. The ethics committee of the Friedrich-Schiller-Universität Jena approved the experimental protocol (Reg. No. FSV 22/086), and the study was conducted in accordance with the guidelines of the Declaration of Helsinki. The pre-registration for this study can be found at:https://osf.io/axstg/?view_only=13f0e8794885499d805e6649670b2f13.

### 2.2 Stimuli

In addition to faces we chose images of chairs as control stimuli, as they show approximate vertical symmetry and a clear upright direction. Both these features seem to be important in norm-based coding of objects (77). As cues we used geometrical shapes of different colours (a green, a blue, and a yellow frame, with similar area, brightness, and saturation levels). All stimuli were presented by Psychtoolbox-3 (MATLAB-based; (78,79)).

We manipulated perceived typicality of faces and chairs as follows. For faces, we followed previous works as regards the stimulus database and the manipulation approach (10,80–82). In detail, we created three-dimensional face stimuli using DI3Dcapture™ (Dimensional Imaging, Glasgow, UK). Each face had been photographed by four cameras simultaneously, and the images were interpolated to create a three-dimensional (3D) object. Using DI3Dview™, we generated caricatures and anti-caricatures (both in shape and texture), by extrapolating each individual veridical 3D file with the morphing tool with respect to a gender-matched average. Finally, because the 3D camera system permits to extract images from various viewing angles, we produced our stimuli by systematically tilting the individual faces according to a set of ten camera angles (see (80) for a more detailed description of this procedure). Based on a previous rating study, we selected 24 face identities (half female and half male). We subjected each face to both caricaturing (shifting both face shape and texture + 0.33 units on the face trajectory, *opposite to* the face norm) and anti-caricaturing (shifting both face shape and texture + 0.33 units on the face trajectory*, in the direction of* the face norm). For each participant, we selected only one caricaturing version per identity (counterbalanced across participants), leading to 12 caricatured (distinctive) and 12 anti-caricatured (typical) identity for each participant. Throughout the manuscript we use the term “typical” to refer to both faces morphed towards the average and to chairs rated as typical. Conversely, the term “distinctive” refers both to faces morphed in the opposite direction with respect to the average and to chairs rated as distinctive. Instead, “anti-caricatured” and “caricatured” refer to the procedure applied to make faces typical and distinctive, respectively. Since we wanted to avoid image-dependent identity perception (i.e., the same picture of Identity 4 is shown throughout the experiment), we produced 10 different images for each identity. We used Adobe Photoshop™ to tilt each face on 10 pre-set angles, resulting in 240 stimuli in total for participant for each task run.

Colour images of chairs were downloaded from the Internet and were subjected to standard pre-processing to ensure similar image size, quality, and a homogeneous background. No chair contains written text or recognizable logos. To make the chairs comparable to face images, we used 10 images per chair, each showing the image on a comparable set of angles. We manipulated typicality by selecting the 12 most typical and the 12 most distinctive chair images, as rated by a separate group of 10 participants (Cronbach’s alpha = 0.991).

Both face and chair stimuli were presented in black and white, to avoid confounding effects due to the wider range of colours of chairs, as compared to faces. See Figure 1 for examples of face and chair stimuli.

**Figure 1.**
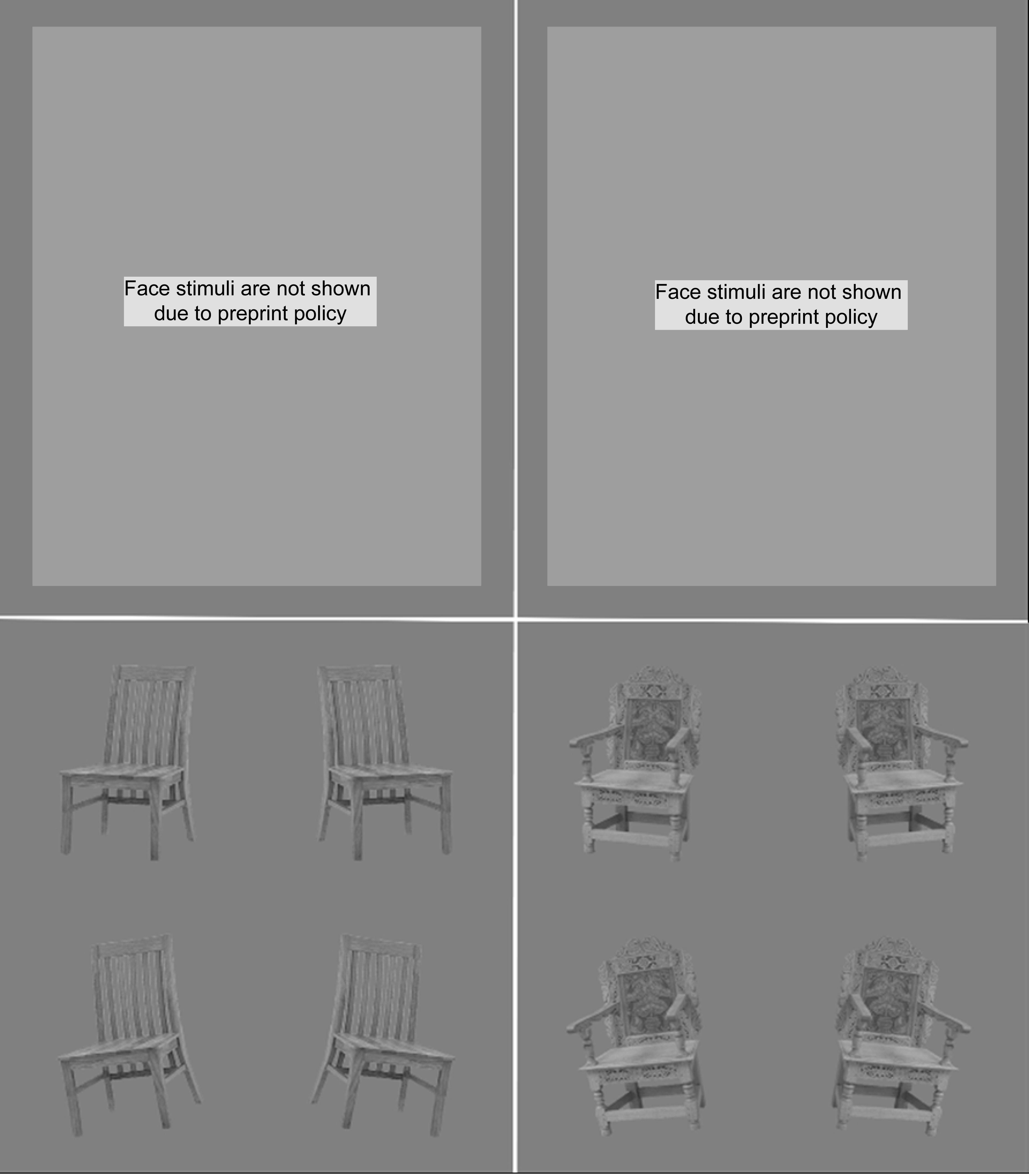
Stimulus examples. We show here the same face identity in its caricatured and anti-caricatured version for comparison. Note that each identity was shown to each participant in either of the two versions only. Left: a typical (anti-caricatured) face and a typical chair. Right: a distinctive (caricatured) face and a distinctive chair.

**Figure 2:**
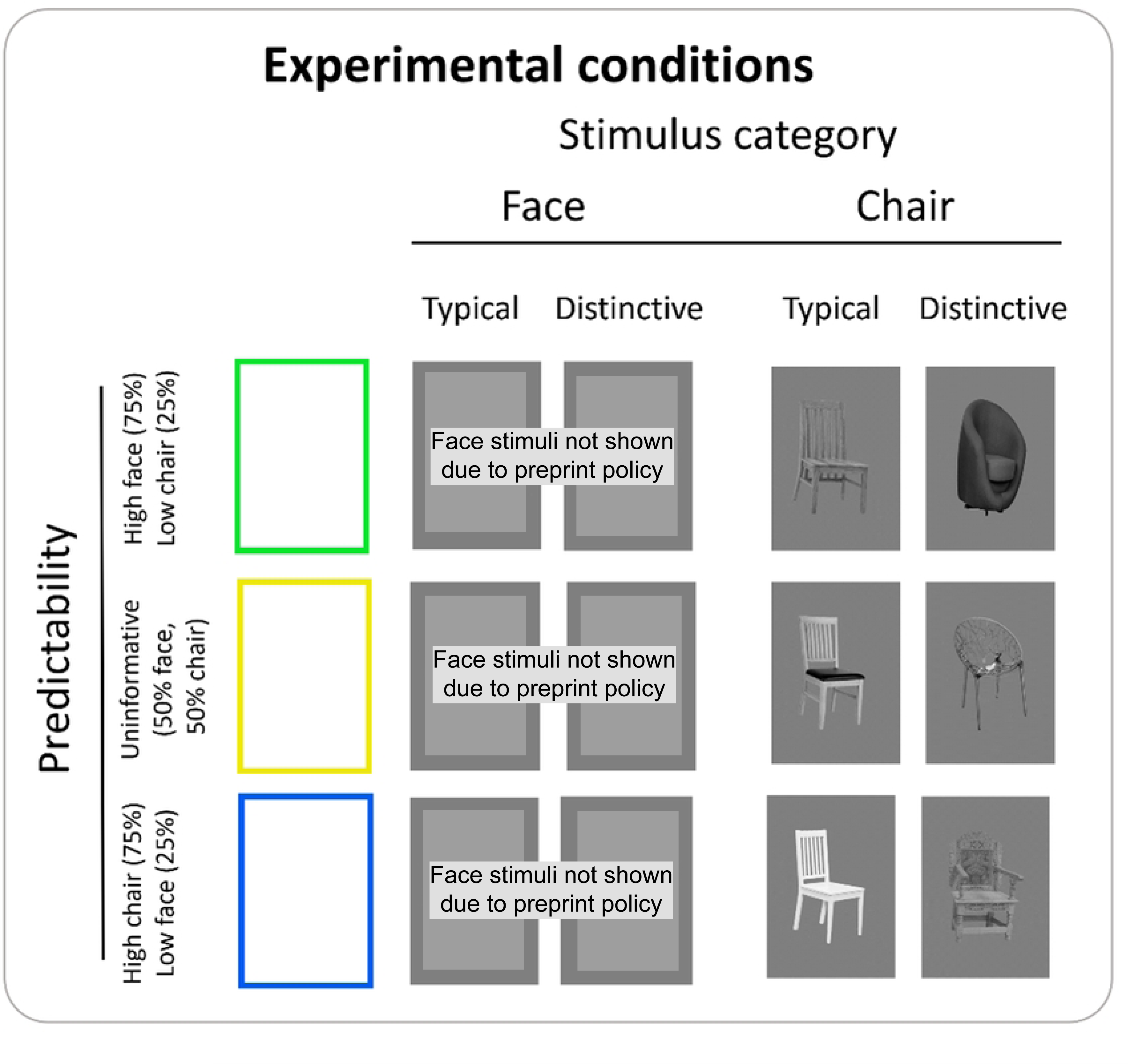
Graphical representation of the experimental conditions.

### 2.3 Experimental Design and Procedure

To manipulate predictability of stimulus category (face or chair), a green frame preceded a face in 75% of the trials, and a chair in the remaining 25% of trials. Conversely, a blue frame preceded a chair in 75% of the trials and a face in the remaining 25%. As suggested by (83) and following (74), we included a neutral, uninformative condition, whereby a yellow frame preceded faces or chairs equally often (50% of the trials). Throughout the task, participants saw 24 identities (12 faces + 12 chairs), in 10 camera angles. Participants performed four task runs in total (two participants performed only three runs due to technical issues). The order of trials within each experimental run was randomized. Each trial started with a fixation cross, which also separated trials (2000-4000 ms, plus a blank if participants responded, to ensure that the duration of one trial is an integer multiple of our TR = 2), followed by a cue (250 ms), and a target stimulus (750 ms). To ensure participants’ attentional focus, in 10% of these trials the stimulus was inverted (but otherwise followed the same cue contingencies as used for upright stimuli), and participants performed an orthogonal detection task of these trials by pressing a key to inverted stimuli with their right index finger on a MRT-compatible keyboard. The experiment included a total of 528 trials, divided in 4 runs of 132 trials each (39 usable, non-target trials per condition in each run). Target trials were excluded from analyses. Participants were informed about the cue contingencies, but we specified that these were not task relevant. We did so with the aim to replicate the design by (74), who instructed their participants in a similar way. Before completing the task, participants were familiarized with the procedure and the contingencies through a short practice session (∼2.5 minutes) outside the scanner.

After completing the fMRI measurement, participants completed a short questionnaire in which they reported the degree to which they experienced any difficulties during scanning, paid attention to the contingencies, and thought the faces and chairs looked typical or distinctive (cf. Supplementary Materials).

### 2.4 Imaging Parameters and Data Analysis

We used a 3-Tesla MR scanner (Siemens MAGNETOM Prisma fit, Erlangen, Germany) with a 20-channel head coil to record BOLD responses to our manipulations. T1-weighted high-resolution 3D anatomical images were acquired with an MP-RAGE sequence (192 slices; TR = 2300 ms; TE = 3.03 s; flip angle = 90°; 1 mm isotropic voxel size). As for the four functional runs, T2*-weighted fMRI-images were collected with a multi-band EPI sequence (MB acceleration factor = 8) under the following parameters: 34 slices; FOV = 204 x 204 mm2; TR = 2000 ms; TE = 30 ms; flip angle = 90°; 2 mm isotropic voxel size. Similarly to a previous study (84), we used SPM12 (Welcome Department of Imaging Neuroscience, London, UK), based on MATLAB version R2020a, to pre-process data.

The experiment was based on an event-related design. Functional images were slice-timed, realigned (the structural image was realigned to a mean image computed from the functional series, and co-registered to structural scans). We then normalized the images to the MNI-152 space, resampled to 2 x 2 x 2 mm resolution, and spatially smoothed using an 8-mm Gaussian kernel. The convolution of a reference hemodynamic response function (HRF) with box cars, representing the onsets and durations of the experimental conditions, was used to define the regressors for a general linear model analysis of the data. For each of the experimental conditions we modelled the HRF, synchronised to the onset of the trial. Low frequency components were excluded from the model using a high-pass filter with 128 s cut-off. Variance which could be explained by the previous scans was excluded using an autoregressive AR(1) model, and movement related variance was accounted for by the spatial parameters resulting from the realignment procedure. The resulting regressors were fitted to the observed functional time series. For the random effects analysis, the contrast estimates entered a simple t-test or an F-test at the second level (see Table S3 for the regressor weights in the whole-brain analysis). Results were thresholded at p < 0.05 (voxelwise, uncorrected), with a cluster correction for multiple comparisons at FWE < 0.05. For visualization, the thresholded t-images were superimposed onto a standard template available on MRIcroGL (https://www.nitrc.org/projects/mricrogl/). We extracted the peaks of maximal activation and their respective anatomical labels using the Anatomy Toolbox implemented in (93) and the Talairach Daemon Client (94,95).

Additionally, participants performed a 6-minutes functional localizer task, so that we could isolate regions of interest (ROIs). Faces, everyday objects, and Fourier-randomized noise images were presented (4 Hz; 230ms exposition time; 20 ms ISI) in blocks of 10 s, interrupted by breaks of 10 s and repeated five times. Each block included 40 images (size: 400 x 400 pixels with a grey background). Participants were instructed to observe the images and maintain their attentional focus on the screen. The fusiform face area (FFA; (85,86)) and the lateral occipital complex (LOC; (87,88)) were isolated. The object-selective area LO (87,89) corresponded to the posterior dorsal portion of the lateral occipital complex (LOC, (90)). We determined the location of the FFA in individual participants by contrasting face and object blocks (face and noise blocks, if the former led to no significant voxels) and established as the local maximum from the t-maps with a threshold of *p*_*FWE*_< 0.05 on the single-subject level. A similar approach was taken to locate the LOC (objects > noise blocks, t-maps with a threshold of *p*_*FWE*_< 0.05), on the single-subject level. We report the individual MNI coordinates in Table S1, and report the average (± SD) here: right FFA (*x*_*M*_= 40.6, *x*_*SD*_ = ± 4.7; *y*_*M*_= −52.9, *y*_*SD*_ = ± 7.7; *z*_*M*_= −18.6, *z*_*SD*_ = ± 3.9), left FFA (*x*_*M*_= −38.9, *x*_*SD*_ = ± 3.5; *y*_*M*_= −52.4, *y*_*SD*_ = ± 6.9; *z*_*M*_= −18.9, *z*_*SD*_ = ± 4.0), right LOC (*x*_*M*_= 40.1, *x*_*SD*_ = ± 4.6; *y*_*M*_= −76.1, *y*_*SD*_ = ± 8.2; *z*_*M*_= −3.6, *z*_*SD*_ = ± 6.4), left LOC (*x*_*M*_= −43.1, *x*_*SD*_ = ± 5.1; *y*_*M*_= −78.3, *y*_*SD*_ = ± 6.6; *z*_*M*_= −3.5, *z*_*SD*_ = ± 5.7). Areas matching anatomical criteria were considered as their appropriate equivalents on the single subject level. A time-series of the mean voxel value within a 4 mm radius sphere around the local peak was extracted from our event related sessions using finite impulse response (FIR) models (91) for each area and participant separately. As for the main task analysis, we convolved our data with a reference HRF using boxcars, to define the regressors for a general linear model using MarsBaR 0.44 toolbox for SPM 12 (92). The peak BOLD values (corresponding to the third TR post-stimulus onset) were extracted from the four event-related runs while trials with upside-down stimuli were excluded from the analysis. We analysed only upright trials using repeated-measures ANOVAs in each ROI with cue-induced Predictability (3, high-face expectation, uninformative, high-chair expectation), Category (2, faces and chairs) and Typicality (2, typical and distinctive) as within-subject factors. We were especially interested in the main effects of typicality (both overall and separate by category), the interaction between category and predictability (replication of (74)’s results) and, critically, the interaction between typicality and predictability. These analyses were performed on JASP (Version 0.16.3). All multiple comparisons of post-hoc tests were corrected with Holm’s method.

## 3. Results

### 3.1 Behavioural Results

Participant’s accuracy was close to ceiling (*M* = 0.99 proportion correct responses, *SD* = 0.01, response time: *M* = 543 ms, *SD* = 77 ms). This suggests that they attended to the task well, and that the task was easy to perform. Due to the small number of inverted trials, we did not perform statistical analysis, but note that numerically, participants were faster for the detection of inverted faces than for inverted chairs, *M* = 525.4 vs. 553.4 ms, respectively. All participants confirmed that experimental faces were similar to those encountered throughout their life. On average participants reported only relatively low attention to cue contingencies (*M* = 0.77 on a scale from 0 to 3; see Table S2 for details) and we detected no effects of predictability on RT (*p* = 0.1692). Note that participants were instructed that cue contingencies were irrelevant to task completion. Finally, the presence of a typicality manipulation was subjectively noticed for chairs more often than for faces, *M* = 20/35 vs. 5/35 participants, respectively. We exploratively investigated behavioural effects of typicality, but found no differences between typical and distinctive stimuli, neither on RTs (*p* = .163) nor in accuracy (F(1, 416) = 1.237, *p* = .267).

### 3.2 Neuroimaging Results

#### 3.2.1 Regions-of-Interest

We found prominent effects of Category and Typicality in all our ROIs (Figure 3). Here we report the average between left and right hemisphere, see the Supplementary Materials the results for the individual ROIs. Considering all participants, on a total of 12600 non-target trials (counted considering all participants together), only 32 were false alarms. Since this number is low, and we did not have reasons to expect these to affect any of our experimental conditions differently, we did not remove them from brain data analysis. In the FFA the main effect of Category was significant 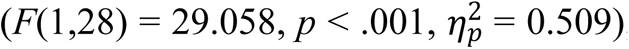, with faces eliciting larger responses than chairs (*p =* .001). The effect of Category was also significant in the LOC 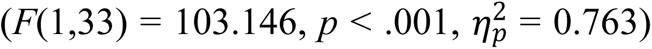, where chairs elicited larger responses than faces. Considering that these regions were selected based on the functional localizer and extensive previous literature, these results are expected, and can be seen as an indicator of data quality. More interesting are the effects of Typicality over the four ROIs (all *p*s < .001), which were at times comparably sizable as those of Category (e.g., right FFA, 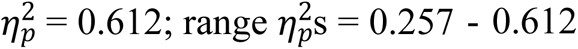). The effect of Typicality was significant and in the same direction even when considering only the stimulus Category for which each ROI is specialized (i.e., faces for FFAs and chairs for LOCs; all *ps*_*Holm*_< .005; see Figure S1). Remarkably, the hypothesized interactions between stimulus Typicality and Predictability failed to reach significance (cf. Table 2 for details). We additionally analysed this interaction when only considering the preferred stimulus category of the ROIs (faces for FFA, chairs for LOC). Again, the interactions did not reach significance, although in the LOC we note a trend (*p* = .081), with distinctive chairs eliciting larger responses than typical chairs especially in the uninformative condition. In line with (74), we found no main effect of Predictability in any ROI. We also could not replicate the interaction between Category and Predictability reported by (74) either, and no other interactions reached significance. We additionally report the Bayes Factor for the main models we tested in the FFAs and the LOCs. In both cases, the model with the highest evidence in favour of the alternative hypothesis was that including the two main effects of Category and Typicality but not their interaction (*BF*_10_ = 1.000 in the FFAs, and *BF*_10_= 1.000 in the LOCs). These values can be interpreted as weak evidence in favour of the described model (96). Additionally, in both cases the model with the strongest evidence in favour of the null hypothesis is that including solely the main effect of Predictability (*BF*_10_ = 5.363^―7^in the FFAs, and *BF*_10_ = 7.147^―13^in the LOCs). The analysis for our model of interest, including the main effects of Category, Typicality and Predictability, plus the interactions Predictability*Typicality, returned strong support in favour of the null hypothesis both in the FFAs (*BF*_10_= 0.005) and in the LOCs (*BF*_10_= 0.028) (96). Finally, the model including Category, Predictability and their interaction in the FFAs, which would correspond to the effect reported in (74) received strong evidence in favour of the null hypothesis (*BF*_10_ = 2.234^―4^). Overall, these analyses are consistent with the effects calculated under a frequentist framework - although the evidence in favour of our two main effects remains weak and point to the absence of our interaction of interest.

**Figure 3:**
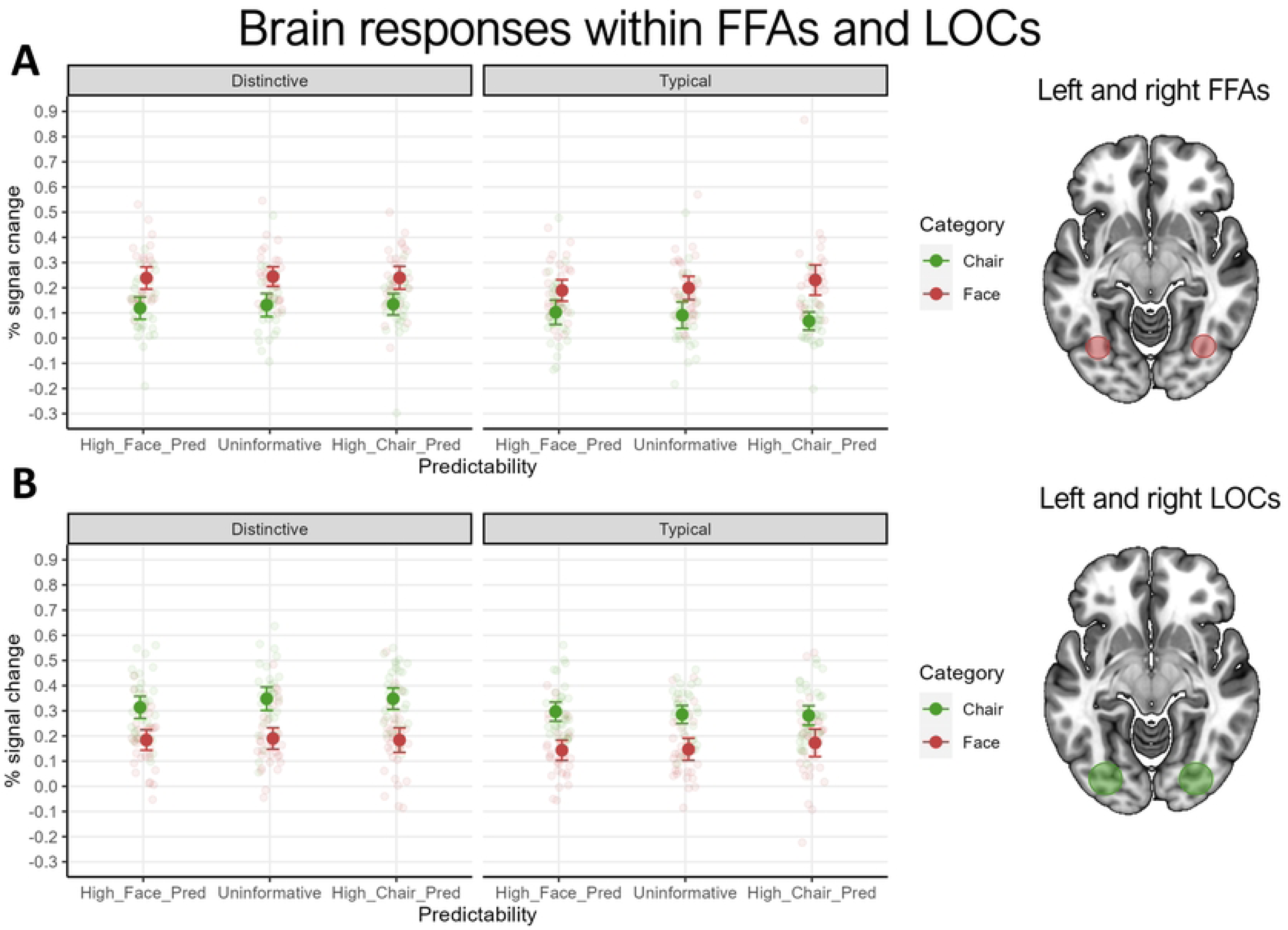
Results of the ROI analysis. Brain responses to typical and distinctive stimuli of the two categories to facilitate the comparison with Figure 3 in (74). High_Face_Pred: cue that highly predicts a face; High_Chair_Pred: cue that highly predicts a chair; Uninformative: cue that predicts either category equally; panel A: brain responses to typical and distinctive stimuli in both left and right FFAs averaged; panel B: brain responses to typical and distinctive stimuli in both left and right LOCs averaged. For a figure depicting brain responses in individual ROIs, see Figure S1. ROIs shown on the right for illustration purposes only, the average locations of the individually defined ROIs are indicated in Section 2.4. Error bars represent 95% confidence intervals.

**Figure 4:**
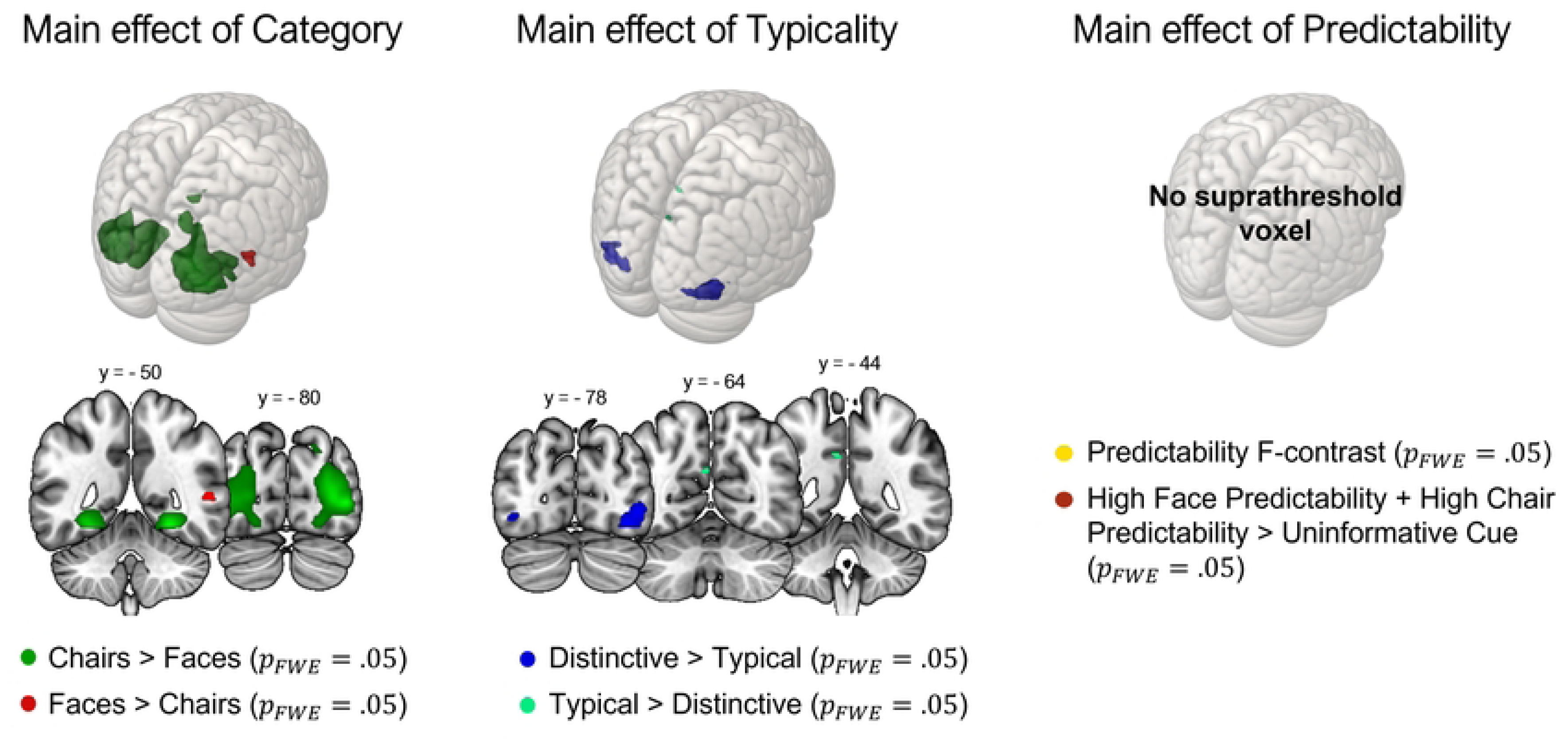
Whole brain results. (MNI standard space). From left to right: directional t-contrasts of brain activation in response to faces > chairs (red) and chairs > faces /green); directional t-contrasts of brain activations in response to distinctive > typical stimuli (dark blue) and typical > distinctive (light green); non-directional F contrast assessing whether any of the predictability levels differs from the others (yellow) and directional t-contrast comparing the two conditions of high predictability to the uninformative condition (dark red). All contrasts are shown at an alpha level of .05, with a family-wise correction for multiple comparisons.

**Table 1:**
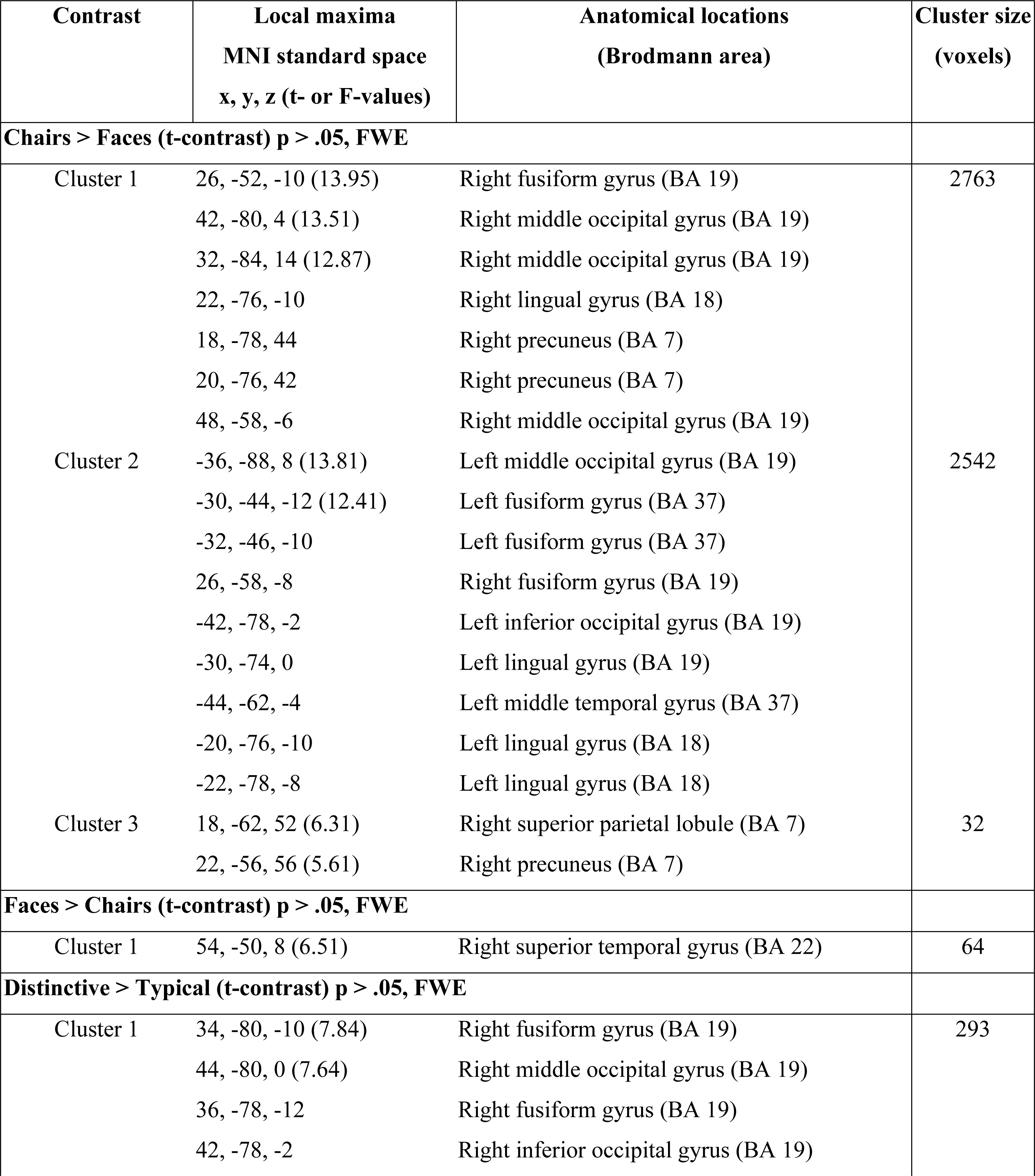
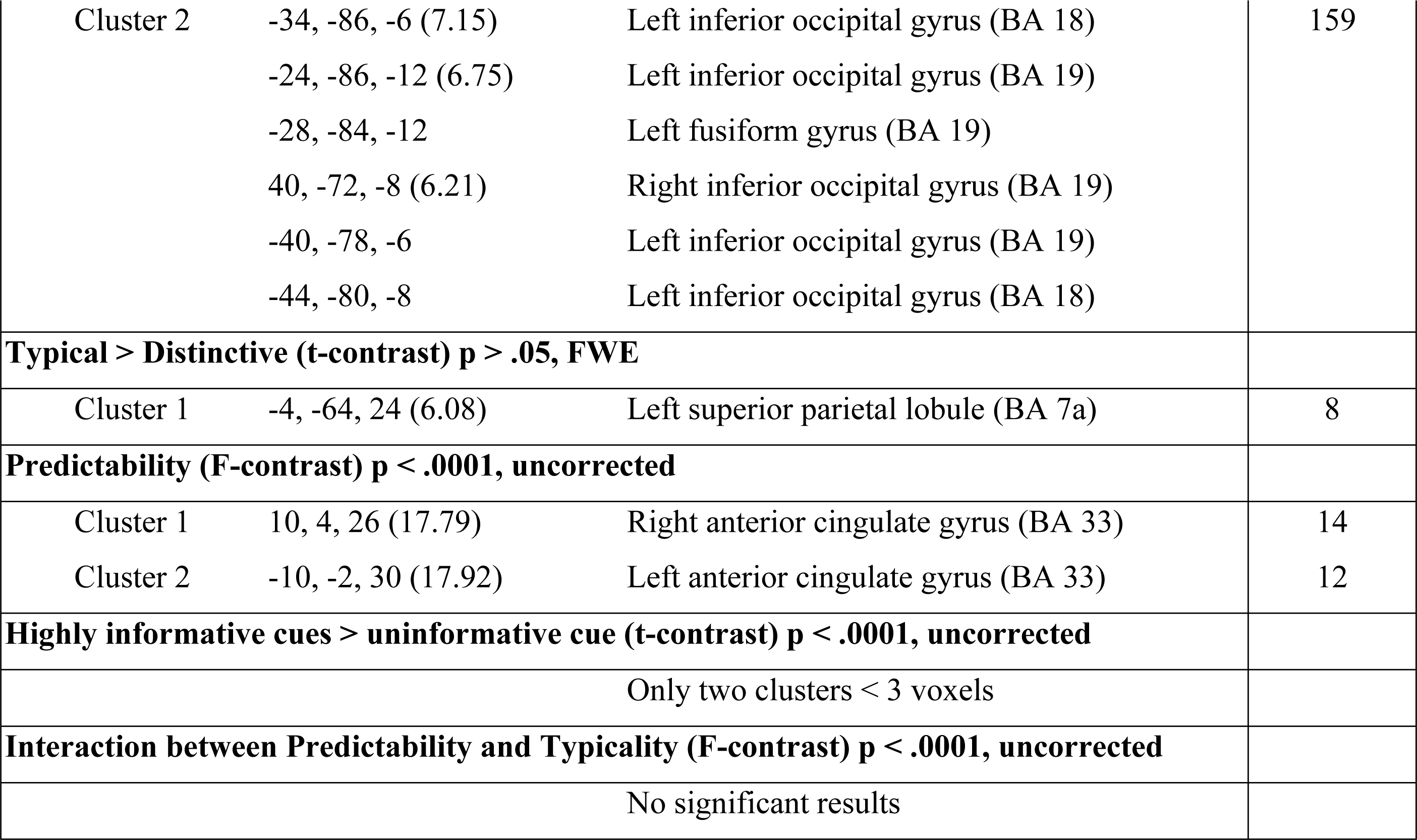
Local maxima of activation for the contrasts of interest at the whole brain level. Note that t- or F-values are provided for the peaks output by SPM12, whereas the other peaks have been located with the Anatomy Toolbox. For them all, the third column reports the anatomical location assigned via the Anatomy Toolbox. The raw output of the Anatomy Toolbox is provided as Supplementary Materials.

**Table 2.**
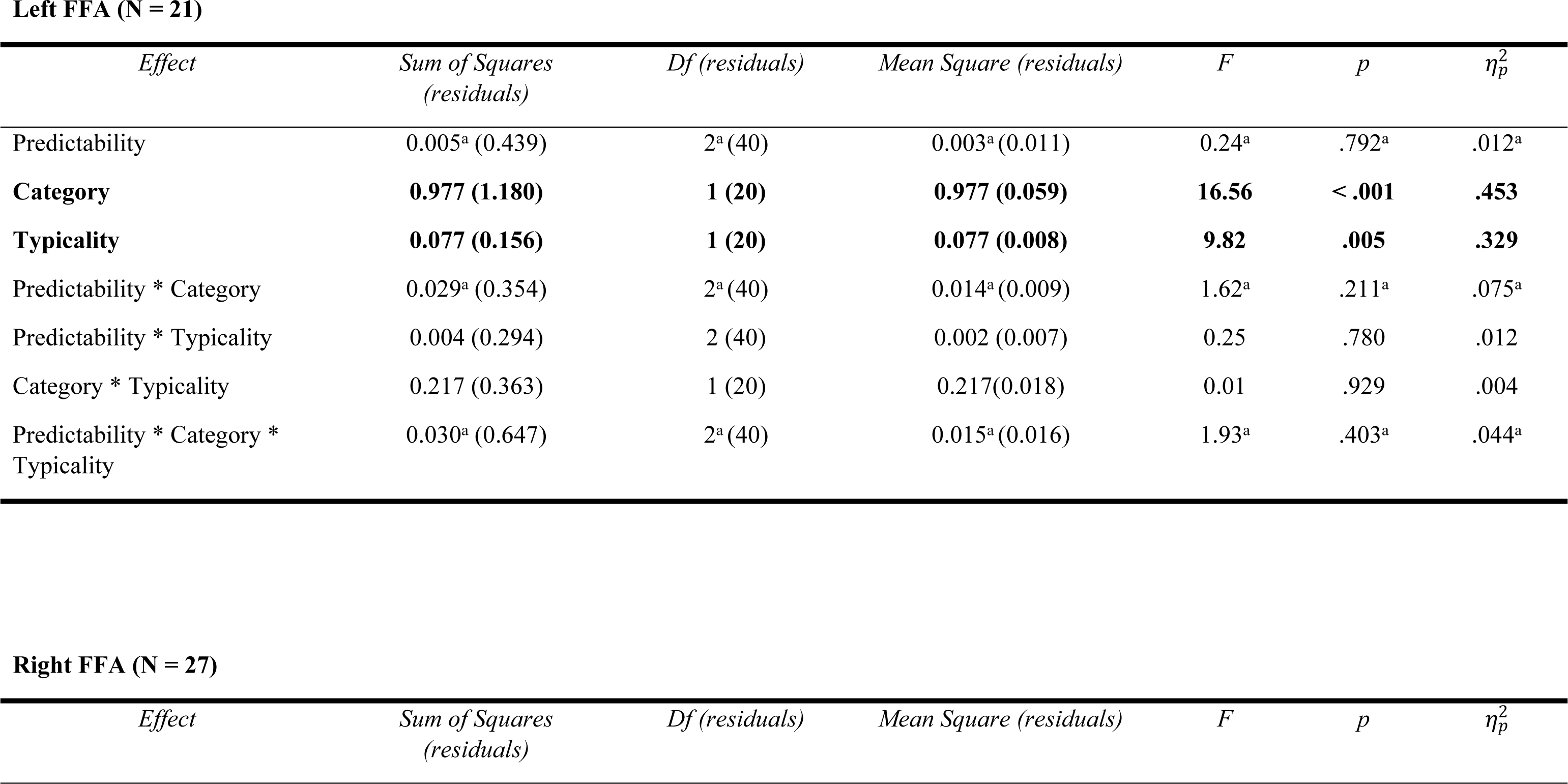
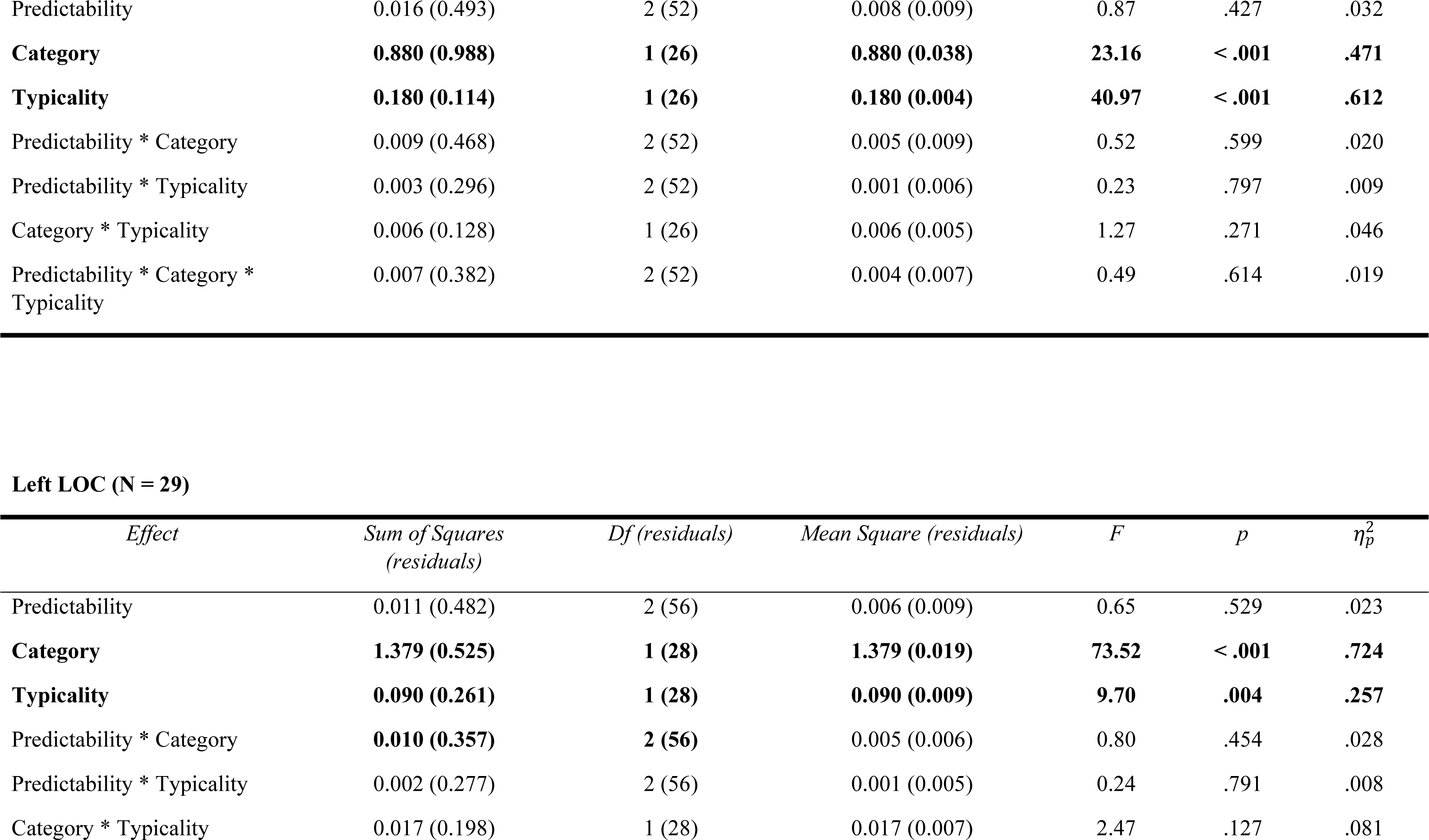
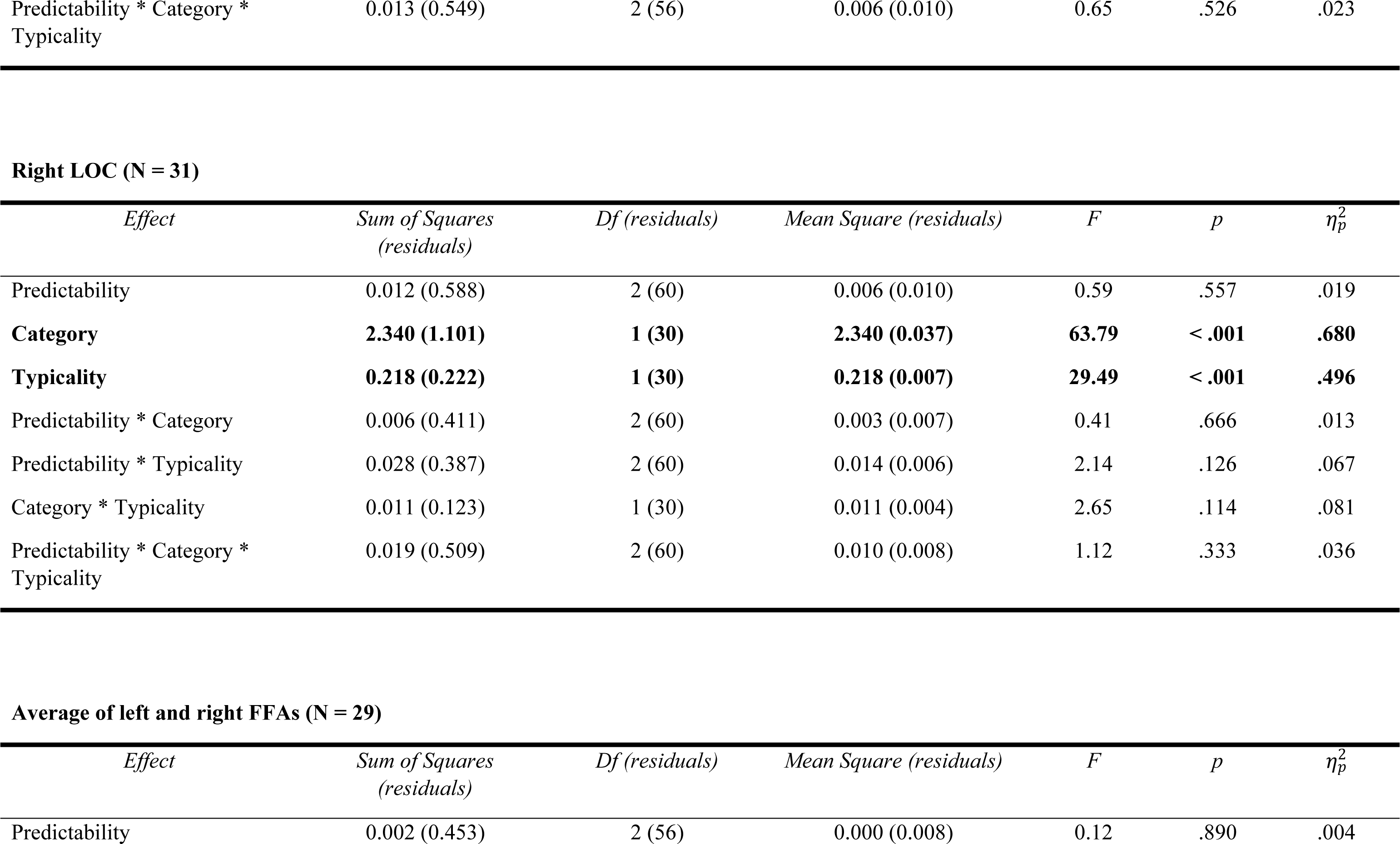
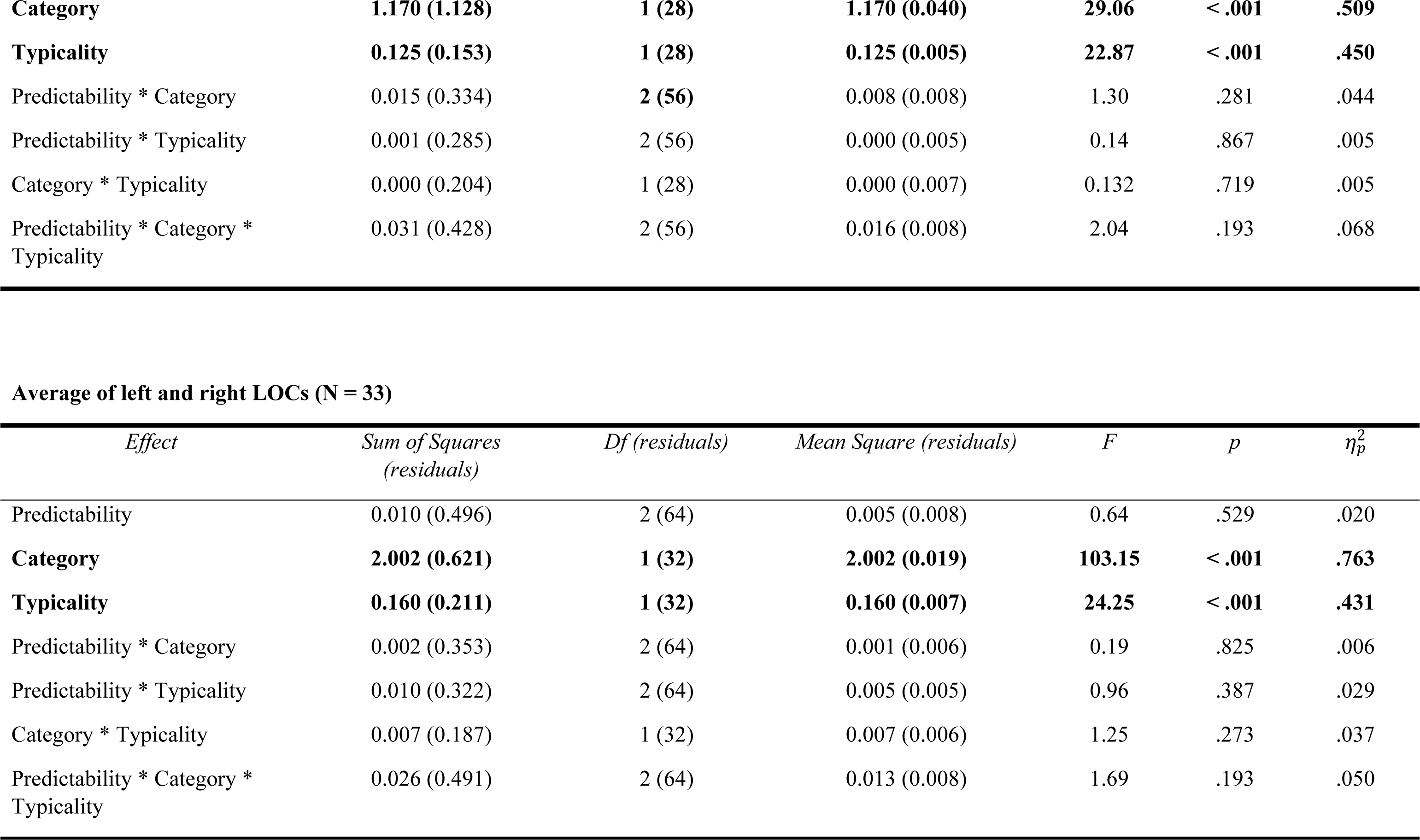
Within-subjects ANOVAs results and contrast weights. Within-subjects ANOVAs with Predictability (3 levels), Category (2 levels) and Typicality (2 levels) as within-subjects factors. Results are presented both in each ROI singularly and in face- and object-responsive regions averaged across hemispheres. Significant effects are boldened. Sum of squares of type III are reported. ^a^ Mauchly’s test of sphericity indicates that the assumption of sphericity is violated (p < .05).

**Left FFA (N = 21)**
| *Effect* | *Sum of Squares*<br>*(residuals)* | *Df (residuals)* | *Mean Square (residuals)* | *F* | *p* | *$\eta_p^2$* |
| --- | --- | --- | --- | --- | --- | --- |
| Predictability | 0.005<sup>a</sup> (0.439) | 2<sup>a</sup> (40) | 0.003<sup>a</sup> (0.011) | 0.24<sup>a</sup> | .792<sup>a</sup> | .012<sup>a</sup> |
| **Category** | **0.977 (1.180)** | **1 (20)** | **0.977 (0.059)** | **16.56** | **< .001** | **.453** |
| **Typicality** | **0.077 (0.156)** | **1 (20)** | **0.077 (0.008)** | **9.82** | **.005** | **.329** |
| Predictability * Category | 0.029<sup>a</sup> (0.354) | 2<sup>a</sup> (40) | 0.014<sup>a</sup> (0.009) | 1.62<sup>a</sup> | .211<sup>a</sup> | .075<sup>a</sup> |
| Predictability * Typicality | 0.004 (0.294) | 2 (40) | 0.002 (0.007) | 0.25 | .780 | .012 |
| Category * Typicality | 0.217 (0.363) | 1 (20) | 0.217(0.018) | 0.01 | .929 | .004 |
| Predictability * Category *<br>Typicality | 0.030<sup>a</sup> (0.647) | 2<sup>a</sup> (40) | 0.015<sup>a</sup> (0.016) | 1.93<sup>a</sup> | .403<sup>a</sup> | .044<sup>a</sup> |

**Right FFA (N = 27)**
| *Effect* | *Sum of Squares*<br>*(residuals)* | *Df (residuals)* | *Mean Square (residuals)* | *F* | *p* | *$\eta_p^2$* |
| --- | --- | --- | --- | --- | --- | --- |
| Predictability | 0.016 (0.493) | 2 (52) | 0.008 (0.009) | 0.87 | .427 | .032 |
| **Category** | **0.880 (0.988)** | **1 (26)** | **0.880 (0.038)** | **23.16** | **< .001** | **.471** |
| **Typicality** | **0.180 (0.114)** | **1 (26)** | **0.180 (0.004)** | **40.97** | **< .001** | **.612** |
| Predictability * Category | 0.009 (0.468) | 2 (52) | 0.005 (0.009) | 0.52 | .599 | .020 |
| Predictability * Typicality | 0.003 (0.296) | 2 (52) | 0.001 (0.006) | 0.23 | .797 | .009 |
| Category * Typicality | 0.006 (0.128) | 1 (26) | 0.006 (0.005) | 1.27 | .271 | .046 |
| Predictability * Category * Typicality | 0.007 (0.382) | 2 (52) | 0.004 (0.007) | 0.49 | .614 | .019 |

Left LOC (N = 29)
| *Effect* | *Sum of Squares (residuals)* | *Df (residuals)* | *Mean Square (residuals)* | *F* | *p* | $\eta_p^2$ |
| --- | --- | --- | --- | --- | --- | --- |
| Predictability | 0.011 (0.482) | 2 (56) | 0.006 (0.009) | 0.65 | .529 | .023 |
| **Category** | **1.379 (0.525)** | **1 (28)** | **1.379 (0.019)** | **73.52** | **< .001** | **.724** |
| **Typicality** | **0.090 (0.261)** | **1 (28)** | **0.090 (0.009)** | **9.70** | **.004** | **.257** |
| Predictability * Category | **0.010 (0.357)** | **2 (56)** | 0.005 (0.006) | 0.80 | .454 | .028 |
| Predictability * Typicality | 0.002 (0.277) | 2 (56) | 0.001 (0.005) | 0.24 | .791 | .008 |
| Category * Typicality | 0.017 (0.198) | 1 (28) | 0.017 (0.007) | 2.47 | .127 | .081 |
| Predictability * Category *<br>Typicality | 0.013 (0.549) | 2 (56) | 0.006 (0.010) | 0.65 | .526 | .023 |

**Right LOC (N = 31)**
| *Effect* | *Sum of Squares*<br>*(residuals)* | *Df (residuals)* | *Mean Square (residuals)* | *F* | *p* | $\eta_p^2$ |
| --- | --- | --- | --- | --- | --- | --- |
| Predictability | 0.012 (0.588) | 2 (60) | 0.006 (0.010) | 0.59 | .557 | .019 |
| **Category** | **2.340 (1.101)** | **1 (30)** | **2.340 (0.037)** | **63.79** | **< .001** | **.680** |
| **Typicality** | **0.218 (0.222)** | **1 (30)** | **0.218 (0.007)** | **29.49** | **< .001** | **.496** |
| Predictability * Category | 0.006 (0.411) | 2 (60) | 0.003 (0.007) | 0.41 | .666 | .013 |
| Predictability * Typicality | 0.028 (0.387) | 2 (60) | 0.014 (0.006) | 2.14 | .126 | .067 |
| Category * Typicality | 0.011 (0.123) | 1 (30) | 0.011 (0.004) | 2.65 | .114 | .081 |
| Predictability * Category *<br>Typicality | 0.019 (0.509) | 2 (60) | 0.010 (0.008) | 1.12 | .333 | .036 |

**Average of left and right FFAs (N = 29)**
| *Effect* | *Sum of Squares*<br>*(residuals)* | *Df (residuals)* | *Mean Square (residuals)* | *F* | *p* | $\eta_p^2$ |
| --- | --- | --- | --- | --- | --- | --- |
| Predictability | 0.002 (0.453) | 2 (56) | 0.000 (0.008) | 0.12 | .890 | .004 |
| **Category** | **1.170 (1.128)** | **1 (28)** | **1.170 (0.040)** | **29.06** | **< .001** | **.509** |
| **Typicality** | **0.125 (0.153)** | **1 (28)** | **0.125 (0.005)** | **22.87** | **< .001** | **.450** |
| Predictability * Category | 0.015 (0.334) | **2 (56)** | 0.008 (0.008) | 1.30 | .281 | .044 |
| Predictability * Typicality | 0.001 (0.285) | 2 (56) | 0.000 (0.005) | 0.14 | .867 | .005 |
| Category * Typicality | 0.000 (0.204) | 1 (28) | 0.000 (0.007) | 0.132 | .719 | .005 |
| Predictability * Category * Typicality | 0.031 (0.428) | 2 (56) | 0.016 (0.008) | 2.04 | .193 | .068 |

**Average of left and right LOCs (N = 33)**
| *Effect* | *Sum of Squares (residuals)* | *Df (residuals)* | *Mean Square (residuals)* | *F* | *p* | *$\eta_p^2$* |
| --- | --- | --- | --- | --- | --- | --- |
| Predictability | 0.010 (0.496) | 2 (64) | 0.005 (0.008) | 0.64 | .529 | .020 |
| **Category** | **2.002 (0.621)** | **1 (32)** | **2.002 (0.019)** | **103.15** | **< .001** | **.763** |
| **Typicality** | **0.160 (0.211)** | **1 (32)** | **0.160 (0.007)** | **24.25** | **< .001** | **.431** |
| Predictability * Category | 0.002 (0.353) | 2 (64) | 0.001 (0.006) | 0.19 | .825 | .006 |
| Predictability * Typicality | 0.010 (0.322) | 2 (64) | 0.005 (0.005) | 0.96 | .387 | .029 |
| Category * Typicality | 0.007 (0.187) | 1 (32) | 0.007 (0.006) | 1.25 | .273 | .037 |
| Predictability * Category * Typicality | 0.026 (0.491) | 2 (64) | 0.013 (0.008) | 1.69 | .193 | .050 |

#### 3.2.2 Whole-brain analyses

We investigated the main effects of Predictability (both by means a non-directional F contrast and a directional t-contrast comparing high predictability conditions to the uninformative one), Typicality and Category. We also investigated the interaction of interest (Predictability * Typicality). The contrast faces > chairs revealed one cluster in the right superior temporal gyrus (rSTG), whereas the opposite contrast returned activation in two large bilateral clusters mainly encompassing the LOCs, the fusiform gyri, V1, V3, V5, extending to the hippocampal cornu ammonis, and to the inferior and superior parietal lobules. The contrast typical > distinctive generated a small cluster in the left superior parietal lobe, while the opposite (distinctive > typical) revealed two symmetrical clusters which include the LOCs, the fusiform gyri and the ventral portions of V2. We report the same effect estimated in faces and chairs separately in Figure S2. The main effect of Predictability resulted in no suprathreshold clusters (p < .05, FWE corrected), and relaxing the threshold to the conventionally accepted level of p < .001 (uncorrected) resulted in two small clusters in the right hemisphere, one in the anterior cingulate gyrus, and the other in the anterior insula. The directional contrast reflecting high predictability conditions compared to the uninformative one revealed no significant results (p < .05, FWE corrected). Setting a more lenient threshold at p < .0001, uncorrected resulted only in a three-voxel cluster in the right anterior cingulate gyrus which we report, but refrain from interpreting. The contrast relative to the interaction between Predictability and Typicality was not significant.

## 4. Discussion

This study investigated combined effects of stimulus typicality and cue-induced predictability. Our results confirm reports of increased responses to distinctive, as compared to typical stimuli for both face and non-face stimuli. In contrast to our hypotheses, we neither found robust effects of predictability, nor an interaction between predictability and typicality.

### 4.1 The Brain is Sensitive to Visual Typicality

We detected stronger brain responses to distinctive, as compared to typical stimuli in all the tested ROIs, which confirms previous findings regarding face typicality encoding in the FFA (28,52,53) and extends them to non-face stimuli and their processing in the LOC. Most participants also seem to have noticed variations of typicality either in faces or chairs, when explicitly asked after the experiment. Critically, the lack of interaction with stimulus category indicated that these regions are sensitive to the distance of the stimulus from the prototype regardless of category, which also appears from our separate analyses of typicality effects for faces and chairs (Figure S2). Overall, these results align both with previous findings about the neural effects of typicality (see Section 1.2) and may represent a neural signal encoding distance from the prototype (also see (10), where increasing distance leads to decreasing P200 amplitude). Note that although this idea is more plausible when considering norm-based models (6,18,26) it also can be reconciled with exemplar-based models of visual perception (22,97,98). In this case, enhanced brain responses to distinctive stimuli might reflect distances of these from the majority of the others (with no need of abstraction). These responses might then represent a “rarity” signal, based on the fact that distinctive stimuli are also less frequent. Finally, reduced responses to typical stimuli might also represent a phenomenon of neural adaptation to statistical regularities associated to that particular category - when a face is perceived, it is more likely to present certain features - in line with predictive accounts of vision (99). However, as discussed below, the extent to which such adaptation concerns category-space distances or rather low-level features remains an open question.

We also found increased brain responses to distinctive, as compared to typical stimuli in two large bilateral clusters including the fusiform gyri, the middle and the inferior occipital gyri in our whole-brain analysis. This aligns with the prediction that typicality processing should occur in regions representing and tying visual features into coherent percepts, such as the LOC (52). It is interesting to compare our univariate whole-brain results with those of a multivariate study reporting that typical stimuli of two different categories are maximally differentiated in the LOC, in which between-category boundaries are thus maximized (29). One interpretation may be that the LOC maximally discriminates between categories by storing prototypical representations, such that its responses to prototype-deviant stimuli can be seen as a potential “surprise” signal. We additionally found a small significant cluster of voxels where activity was larger for typical when compared to distinctive stimuli, corresponding to the inferior parietal lobule (IPL), which is reminiscent of a cluster identified by (29) in a searchlight analysis. In their study, neural patterns in response to distinctive (atypical) exemplars were more differentiated in the caudal IPL. (29) speculate that this area facilitates recognition of atypical items, achieved by relating these items to the respective category ((29), pp. 175-176). Therefore, stronger activation to typical stimuli within the same area might reflect the activation of prototypical representations necessary to establish the category to which the item belongs. We also note that, while typicality information consolidated through the years (like that to faces and common objects) seems to take place in the visual stream and the IPL, other regions in the frontal lobe and the hippocampus might be involved in learning new prototypes, from entirely new visual categories (24). The conceptual separation between *structural* and *functional* typicality (1) is thus mirrored at the neural level.

As a cautionary note, the activation differences between typical and distinctive stimuli may reflect differences in low-level features, rather than differences in the location within a hypothetical category space. While it is difficult to determine the extent to which this is the case in the current study and especially in the case of chair images, we note that we equalized luminance and spatial frequency distribution in the case of faces, for which we still find typicality effects in our ROI analysis. Moreover, (52) found no correlation between a neural measure of physical typicality (height and angle of certain stimuli parts within the same category) and another, based on subjective typicality (ratings). Although we acknowledge that the stimuli used in (52) were more controlled in terms of low-level features and we note that our typicality effects were larger for the less controlled chair stimuli than for faces, these considerations indicate that typicality effects in our study may indeed be driven by category-space structure and not only low-level features.

### 4.2 Failure to Produce Cue-induced Predictability Effects

An important finding is that we detected no main effect of cue-induced predictability in the brain. This likely occurred due to the irrelevance of cue-stimulus contingencies during the task. In fact, while a previous study with a similar cueing design reported lower brain responses in ventral visual areas to strongly expected stimuli (100), (74) also did not find the main effect of cue-based predictability in the fusiform face and parahippocampal place areas. An explanation of the controversial results might be that, in our case and in (74), participants were told explicitly that the cues were task irrelevant, and this corresponds with subjective reports of low attention to cue contingencies in the present study. By contrast, in the study above, participants were explicitly encouraged to learn the cue-stimulus associations and pay attention to them. Moreover, a study where cueing was used for pain conditioning reports activations in the periaqueductal gray when contrasting cues anticipating uncertain pain intensity to those anticipating certain pain intensity (101). Critically, (101) also instructed participants to pay attention to the cues. Since attention enhances overall cortical responsiveness as well as predictability effects (102,103), the lack of predictability effects may be a consequence of the lack of such instructions. However, (104) still found predictability effects in ventral visual areas, despite instructing the participants that the contingencies were not task relevant, similarly to our study and to that of (74). Another hypothesis is that, as (105) suggest, predictability effects in cueing paradigms depend on the presence of a neutral condition (i.e., when a cue is uninformative about the occurrence of stimulus type). Whereas (104), (106) and (101) only included expected and unexpected conditions, our study and that by (74) also included a neutral condition. Moreover, a recent study using uni- and multi-variate analytic approaches on EEG data from a large sample during a cueing paradigm with contingencies similar to ours found no effects of predictability (75). Notably, in (75) participants were intensively and successfully trained to learn the cue-stimulus associations, so the lack of effects cannot be ascribed to lack of attention or failure to use the cues. However, even the idea that the effect can only be found in the absence of a neutral condition is debatable, as (100) report predictability effects face- and object-sensitive regions in the presence of an uninformative condition similar to ours. Similarly, evidence is mixed even regarding whether participants develop predictions about the single stimulus or, rather, about a stimulus category (105). In sum, while it seems to be better to ensure active learning and use of cues, and the effect might be larger in the absence of a neutral condition, cueing designs appear to be less robust as previously thought in producing predictability effects. Ideally, a highly-powered, multi-site and multi-method study should systematically assess the impact of task instructions, attention, cue-stimulus contingencies, and category vs. stimulus learning (see further recommendations in (105)).

Another important finding is that, in contrast to what (74) reported, we failed to detect an interaction between predictability and category, effect on which we based our power analysis. This discrepancy between our results and that of (74), using nearly identical paradigms and instructions, suggests that this effect might not necessarily be robust across smaller differences between the studies (including, for instance, the specific stimulus set used). When inspecting responses for typical and distinctive stimuli separately (Figure 3), we observed a pattern resembling the interaction found by (74), but for typical stimuli only. Although the three-way interaction between category, typicality and predictability was not significant, this pattern suggests that the highly typical faces and buildings used by (74) may have produced enhanced category processing despite the lower number of participants in their study.

### 4.3 Future Investigations on the Interaction between Typicality and Predictability

Critical to our original research question, we detected no interaction between stimulus typicality and predictability, neither in our ROIs, nor at the whole-brain level. We neither found any three-way interaction between typicality, category, and predictability at the ROI level (see Table 1). Our main explanation for the lack of such interactions is the lack of predictability effects the present study, discussed in Section 4.2. An alternative possibility is that, in our manipulations, typicality is formed through years of exposure to stimuli (corresponding to what (1) called “structural typicality”). In contrast, predictability was only induced by verbal instructions, and via prior learning of cue-stimulus associations bearing no semantic values in themselves, and was not task-relevant. The fact that we found strong effects of typicality – whereas predictability effects were absent in the regions of interest and in the rest of the brain – suggests that the two manipulations did not affect neural processing comparably (also see Section 4.3). Finally, we cannot completely exclude the idea that typicality simply does not modulate predictability effects. This would imply a theoretical separation between typicality- and predictability-based neural processing advantages. While predictable stimuli invariably produce smaller prediction error signals (107,108), lower responses to typical stimuli might reflect mechanisms such as neural sharpening (109) or facilitation (110) to commonly seen features.

However, at present these results highlight that cueing designs might not lead to highly robust neural effects, and this affects our ability to make conclusive statements on the presence of interactive vs. additive effects between the two variables, and on their respective theoretical implications. As stated above, further research is needed to obtain a robust cueing paradigm. Afterwards, future studies may test whether typicality and predictability rely on similar or related neural mechanisms by using novel stimulus categories (e.g., (24)), another, different predictability-inducing paradigm (see (83) for suggestions) and compare different models of neural responses (74). Such experiments would allow us to i) manipulate predictability and typicality formed at the same time, ii) detect predictability effects that are strong enough to open the possibility for an interaction (e.g., (107,111)), and iii) establish whether the neural mechanisms driving typicality and predictability are the same.

### 4.4 Category Effects

Finally, our ROI analysis revealed clear differences in the effects of stimulus category between the FFA and the LOC, with stronger responses to faces and to chairs in the two respective areas. In addition, we observed activations in object-responsive regions in the ventral visual stream for chairs (89,90). By contrast, the whole-brain responses to faces did not reveal any regions of the core face processing network (112,113). Instead, faces elicited stronger responses in the rSTG. Typically, studies report face responses in the right inferior/middle temporal gyri (see convergence results in (114) and (113)) and in the superior temporal sulcus (reviewed by (115)), whereas the STG is rarely reported in face research. However, some studies report increased responses in the rSTG to other people’s faces as compared to one’s own (116), and responses to the opposite contrast in its left homotopic area (117,118). In our task, participants saw faces that looked like people their age, that they could encounter in real life, so they might have recruited self-other distinction processes more in response to faces, as compared to chairs.

## 5. Conclusions

We found that distinctive stimuli elicited larger responses in face- and object-responsive regions of the ventral visual stream in ROI analyses, and spatially extended effects encompassing these regions when considering whole-brain analyses. The stronger brain responses to distinctive stimuli are in line with the idea that stimuli that are distant from the prototype (or general tendency of a given category) require more neural resources, and thus produce increased signals. These could be interpreted as a signal of distance from the prototype in the context of prototype-based models, or a rarity signal in the context of exemplar-based models. The present typicality effects seemed to be independent of cue-induced predictability. However, we failed at producing predictability effects with our cueing paradigm and, importantly, we did not replicate the interaction between predictability and category reported by (74). Thus, our research question on the interaction between typicality and predictability remains open, and we provide here some directions for future studies.

## Supporting information

Anatomy Toolbox output

## Acknowledgements

The authors gratefully acknowledge Richard Jahn and Daniel Güllmar for their organizational support during data collection, Julian Kauk for his expert support with data analysis, and our experimental participants for contributing their data to this study. Chenglin Li was supported by the China Scholarship Council (CSC) scholarship (201808330399) during the study. Open Access funding enabled and organized by Projekt DEAL.

## Supplementary captions

**Table S1. Summary of post-experimental survey.** Synopsis of the post-experimental survey completed by participants Each participant was first asked about whether they experienced any form of discomfort during the procedure, and eventually which ones. Then they were asked to indicate to which degree they found themselves paying attention to cue-category contingencies during the scanning session. Their verbal reports (e.g., “never”, not really”, “at the beginning only”, “sometimes”, etc.) were then transferred to a scale ranging from 0 (= “not at all”) to 3 (= “very frequently/always”). No participant reported a score of 3. Participants were then asked whether the faces and the chairs shown during the tasks look ordinary to them or whether they had something distinctive. Their responses were coded as “yes” if they made observations about differential typicality within that category (e.g., “some of the chairs were quite special”), and with “no” if all items of that category appeared of equal typicality to them (e.g., “all faces looked pretty normal to me, none stood out”). As shown in the total proportions of “yes” responses, typicality manipulations were detected much more often in chairs than faces.

**Table S2. ROI locations in individual participants.** MNI coordinates of activation peaks in individual participants during functional localizer run. Right and left fusiform faces areas (FFAs) were localized using the contrast Faces > Objects (or Faces > Noise when the first did not reveal any peak in the approximate area). Right and left lateral occipital complexes (LOCs) were localized using the contrast Objects > Noise. Stars (*) indicate that the coordinate was retrieved when changing the threshold from p_FWE<.05 to p_uncorrected<.001. In a few cases no peaks were found for a particular area. When comparing this table to Fig. S1 and Fig. 3, note that even though we were able to locate these ROIs in nearly all participants, at times the hemodynamic function extracted at the respective location was not of sufficient quality, thus data were excluded from the analysis reported in the main text.

**Table S3. Contrast weights for the whole-brain analysis.** Conditions are abbreviated as follows: H = high predictability cue (75% contingencies); M = medium/uninformative cue (50% contingencies); L = low predictability cue (25 % contingencies); F = face stimulus; C = chair stimulus; T = typical stimulus; D = distinctive stimulus. Note that weights are set to 0 for target trials and for the six nuisance regressors.

**Figure S1. ANOVA results in individual ROIs.**

**Figure S2: Whole-brain typicality effects for faces and chairs separately.** Whole-brain effects of typicality in faces and chairs separately (directional t-contrasts). Results for the contrast distinctive chairs > typical chairs are thresholded at an alpha level of .05 and a family-wise error correction for multiple comparison (FWE, p < .05). The other contrasts are thresholded at an alpha level of .001 and no correction (uncorrected, p < .001). The ink transparency in the legend visually conveys the different thresholding conservativeness of the results. This figure shows that the effects of distinctiveness (distinctive > typical) tend to occur in overlapping regions for both stimulus types (note that, with an uncorrected threshold, the clusters related to chair distinctiveness overlap with those related to face distinctiveness). Conversely, typicality effects seem to be differently distributed and to be more prominent for chairs.

