## Supplementary material for "Investigating the neural effects of typicality and predictability for face and object stimuli": Anatomy Toolbox output

**Raw output of the Anatomy Toolbox**

https://www.fz-juelich.de/en/inm/inm-7/resources/jubrain-anatomy-toolbox

**t-contrast Chairs > Faces ($p_{FWE}$= .05)**

Image: MAIN_t_Chairs-Faces_FWE05 [u=0.0, k=0]

Cluster 1 2763 vox +26 -52 -10

Assignment based on Maximum Probability Map Percent of Cluster volume in Area Percent of Area activated by Cluster

| | | |
|---|---|---|
| Area hOc4la | 9.9 | 16.0 |
| Area hOc4lp | 9.6 | 18.4 |
| Area hOc4v [V4(v)] | 6.8 | 13.7 |
| Area hIP7 (IPS) | 5.6 | 31.7 |
| Area hIP4 (IPS) | 4.4 | 29.7 |
| Area FG3 | 3.5 | 11.4 |
| Area FG1 | 2.6 | 12.2 |
| Area hOc3v [V3v] | 2.4 | 4.0 |
| Area hIP8 (IPS) | 2.3 | 10.5 |
| Area hOc4d [V3A] | 1.4 | 3.9 |
| Area hOc3d [V3d] | 0.8 | 1.7 |
| Area hPO1 (IPS) | 0.7 | 2.9 |
| CA1 (Hippocampus) | 0.7 | 1.9 |
| Area PGp (IPL) | 0.5 | 0.8 |
| Area hIP5 (IPS) | 0.2 | 1.0 |
| Area hOc1 [V1] | 0.2 | 0.2 |
| Area hOc5 [V5/MT] | 0.1 | 1.4 |
| Area 7P (SPL) | 0.0 | 0.1 |
| Subiculum | 0.0 | 0.1 |

| | | |
|---|---|---|
| DG (Hippocampus) | 0.0 | 0.0 |

| Probability exceedance by Area | Mean probability at Cluster location (%) | Mean probability across PMap (%) | Ratio | 95% CI | 95% CI |
|---|---|---|---|---|---|
| Area hOc4la | 37.5 | 30.4 | 1.23 | 1.20 | 1.26 |
| Area hOc4lp | 28.1 | 25.6 | 1.10 | 1.07 | 1.12 |
| Area hOc4v [V4(v)] | 29.7 | 25.3 | 1.17 | 1.14 | 1.19 |
| Area hIP7 (IPS) | 27.0 | 17.4 | 1.55 | 1.50 | 1.60 |
| Area FG3 | 24.2 | 23.6 | 1.03 | 0.99 | 1.06 |
| Area hIP4 (IPS) | 18.1 | 17.0 | 1.07 | 1.02 | 1.10 |
| Area hOc3v [V3v] | 21.8 | 25.2 | 0.87 | 0.84 | 0.89 |
| Area FG1 | 17.7 | 18.9 | 0.94 | 0.90 | 0.97 |
| Area hIP8 (IPS) | 23.6 | 18.9 | 1.25 | 1.20 | 1.29 |
| Area hOc4d [V3A] | 14.0 | 20.1 | 0.69 | 0.67 | 0.72 |
| Area hPO1 (IPS) | 9.1 | 19.2 | 0.48 | 0.45 | 0.50 |
| Area hOc3d [V3d] | 10.1 | 21.3 | 0.48 | 0.46 | 0.50 |
| Area PGp (IPL) | 11.4 | 25.1 | 0.45 | 0.43 | 0.48 |
| CA1 (Hippocampus) | 22.1 | 23.5 | 0.94 | 0.88 | 1.00 |
| Area hIP5 (IPS) | 12.0 | 20.9 | 0.57 | 0.53 | 0.61 |
| Area hOc1 [V1] | 10.0 | 34.8 | 0.29 | 0.26 | 0.31 |
| Area hOc5 [V5/MT] | 7.3 | 15.0 | 0.49 | 0.45 | 0.53 |
| Area hOc2 [V2] | 2.6 | 20.6 | 0.13 | 0.12 | 0.14 |
| Area 7P (SPL) | 9.2 | 22.4 | 0.41 | 0.38 | 0.45 |
| DG (Hippocampus) | 8.2 | 16.2 | 0.51 | 0.44 | 0.59 |
| Subiculum | 8.2 | 21.4 | 0.38 | 0.32 | 0.44 |
| Area FG2 | 2.7 | 20.1 | 0.14 | 0.12 | 0.15 |
| Area hIP6 (IPS) | 6.7 | 19.1 | 0.35 | 0.28 | 0.42 |
| Area FG4 | 2.0 | 24.1 | 0.08 | 0.07 | 0.09 |
| Area 7A (SPL) | 2.7 | 24.2 | 0.11 | 0.10 | 0.12 |
| Area hOc6 [V6] | 1.7 | 20.1 | 0.08 | 0.07 | 0.10 |

| Top probabilites at peak voxels | Union across peaks |
|---|---|
| Area hOc4la | 0.6 |
| Area hOc4lp | 0.6 |
| Area hOc4v [V4(v)] | 0.5 |
| Area hOc3v [V3v] | 0.5 |
| Area hIP8 (IPS) | 0.5 |
| Area hPO1 (IPS) | 0.3 |
| Area hIP4 (IPS) | 0.3 |
| Area 7P (SPL) | 0.1 |
| Area FG3 | 0.1 |
| Area hOc2 [V2] | 0.1 |
| Area hIP7 (IPS) | 0.0 |
| Area hIP5 (IPS) | 0.0 |
| Area 7A (SPL) | 0.0 |
| Area FG2 | 0.0 |
| Area hOc5 [V5/MT] | 0.0 |

| Probabilities at local maxima | +26.0 / -52.0 / -10.0 (Height: 14) | +42.0 / -80.0 / +4.0 (Height: 14) | +32.0 / -84.0 / +14.0 (Height: 13) | +22.0 / -76.0 / -10.0 (Height: 8.9) | +18.0 / -78.0 / +44.0 (Height: 7.2) | +20.0 / -76.0 / +42.0 (Height: 7.2) | +48.0 / -58.0 / -6.0 (Height: 6.4) |
|---|---|---|---|---|---|---|---|
| Area hOc4la | 0.0 | 62.7 | 2.0 | 0.0 | 0.0 | 0.0 | 0.0 |
| Area hOc4lp | 0.0 | 37.2 | 32.1 | 0.0 | 0.0 | 0.0 | 0.0 |
| Area hOc4v [V4(v)] | 0.0 | 0.0 | 0.0 | 47.6 | 0.0 | 0.0 | 0.0 |
| Area hOc3v [V3v] | 0.0 | 0.0 | 0.0 | 47.3 | 0.0 | 0.0 | 0.0 |
| Area hIP8 (IPS) | 0.0 | 0.0 | 0.0 | 0.0 | 19.5 | 32.1 | 0.0 |
| Area hPO1 (IPS) | 0.0 | 0.0 | 0.0 | 0.0 | 6.0 | 24.3 | 0.0 |
| Area hIP4 (IPS) | 0.0 | 0.0 | 28.3 | 0.0 | 0.0 | 0.0 | 0.0 |
| Area 7P (SPL) | 0.0 | 0.0 | 0.0 | 0.0 | 11.5 | 3.3 | 0.0 |
| Area FG3 | 11.4 | 0.0 | 0.0 | 0.0 | 0.0 | 0.0 | 0.0 |
| Area hOc2 [V2] | 0.0 | 0.0 | 0.0 | 5.1 | 0.0 | 0.0 | 0.0 |

| | | | | | | | |
|---|---|---|---|---|---|---|---|
| Area hIP7 (IPS) | 0.0 | 0.0 | 4.5 | 0.0 | 0.0 | 0.0 | 0.0 |
| Area hIP5 (IPS) | 0.0 | 0.0 | 0.0 | 0.0 | 0.0 | 3.2 | 0.0 |
| Area 7A (SPL) | 0.0 | 0.0 | 0.0 | 0.0 | 0.3 | 0.0 | 0.0 |
| Area FG2 | 0.0 | 0.0 | 0.0 | 0.0 | 0.0 | 0.0 | 0.2 |
| Area hOc5 [V5/MT] | 0.0 | 0.1 | 0.0 | 0.0 | 0.0 | 0.0 | 0.0 |

Area hIP6 (IPS)

Investigating the neural effects of typicality and predictability for face and object stimuli

M. Richter, K. Amunts, H. Mohlberg, S. Bludau, S.B. Eickhoff, K. Zilles, S. Caspers. Cytoarchitectonic segregation of human posterior intraparietal and adjacent parieto-occipital sulcus and ist relation to visuomotor and cognitive functions. Cereb Cortex, in press, 2018

Area hPO1 (IPS)

M. Richter, K. Amunts, H. Mohlberg, S. Bludau, S.B. Eickhoff, K. Zilles, S. Caspers. Cytoarchitectonic segregation of human posterior intraparietal and adjacent parieto-occipital sulcus and ist relation to visuomotor and cognitive functions. Cereb Cortex, in press, 2018

Cluster 2 2542 vox -36 -88 +8

Assignment based on Maximum Probability Map Percent of Cluster volume in Area Percent of Area activated by Cluster

Area hOc4lp 10.3 18.2

Area hOc4la 7.4 10.9

Area hOc4v [V4(v)] 7.2 13.5

Area FG3 6.1 18.1

Area hOc4d [V3A] 3.2 8.5

Area hIP7 (IPS) 3.1 16.1

Area FG1 2.9 12.4

Area hOc5 [V5/MT] 2.5 36.9

Area hIP4 (IPS) 1.9 12.0

Area hOc3v [V3v] 1.8 2.8

Area hOc1 [V1] 1.1 0.9

Area hOc3d [V3d] 1.0 2.0

CA1 (Hippocampus) 0.9 2.3

Area FG2 0.2 0.6

Subiculum 0.2 0.6

Area hIP5 (IPS) 0.1 0.3

Area hOc2 [V2] 0.1 0.1

Area hIP8 (IPS) 0.1 0.3

Area FG4 0.0 0.1

| | | |
|---|---|---|
| Area hPO1 (IPS) | 0.0 | 0.0 |

| Probability exceedance by Area | Mean probability at Cluster location (%) | Mean probability across PMap (%) | Ratio | 95% CI | 95% CI |
|---|---|---|---|---|---|
| Area hOc4lp | 30.1 | 25.6 | 1.18 | 1.15 | 1.20 |
| Area hOc4la | 28.6 | 30.4 | 0.94 | 0.92 | 0.96 |
| Area FG3 | 29.3 | 23.6 | 1.24 | 1.21 | 1.28 |
| Area hOc4v [V4(v)] | 31.1 | 25.3 | 1.23 | 1.20 | 1.26 |
| Area hOc3v [V3v] | 18.5 | 25.2 | 0.73 | 0.71 | 0.75 |
| Area FG1 | 18.6 | 18.9 | 0.98 | 0.95 | 1.03 |
| Area hOc4d [V3A] | 18.4 | 20.1 | 0.92 | 0.88 | 0.95 |
| Area hIP4 (IPS) | 22.3 | 17.0 | 1.32 | 1.27 | 1.36 |
| Area hOc5 [V5/MT] | 24.3 | 15.0 | 1.62 | 1.55 | 1.70 |
| Area hIP7 (IPS) | 21.3 | 17.4 | 1.23 | 1.17 | 1.29 |
| Area hOc3d [V3d] | 14.9 | 21.3 | 0.70 | 0.67 | 0.73 |
| Area hOc1 [V1] | 14.7 | 34.8 | 0.42 | 0.40 | 0.45 |
| CA1 (Hippocampus) | 21.7 | 23.5 | 0.92 | 0.87 | 0.98 |
| Area FG2 | 8.0 | 20.1 | 0.40 | 0.38 | 0.42 |
| Area hOc2 [V2] | 4.7 | 20.6 | 0.23 | 0.21 | 0.24 |
| Area FG4 | 6.0 | 24.1 | 0.25 | 0.23 | 0.27 |
| Subiculum | 14.1 | 21.4 | 0.66 | 0.59 | 0.74 |
| Area hIP8 (IPS) | 9.0 | 18.9 | 0.48 | 0.42 | 0.53 |
| Area hPO1 (IPS) | 5.8 | 19.2 | 0.30 | 0.28 | 0.33 |
| DG (Hippocampus) | 8.6 | 16.2 | 0.53 | 0.46 | 0.60 |
| Area hIP5 (IPS) | 9.8 | 20.9 | 0.47 | 0.38 | 0.57 |
| Area PGp (IPL) | 6.2 | 25.1 | 0.25 | 0.19 | 0.30 |
| Area hOc6 [V6] | 1.6 | 20.1 | 0.08 | 0.05 | 0.11 |

| Top probabilites at peak voxels | Union across peaks |
|---|---|
| Area hOc4v [V4(v)] | 0.9 |
| Area FG3 | 0.9 |

| | |
|---|---|
| Area hOc4lp | 0.8 |
| Area hOc3v [V3v] | 0.6 |
| Area hOc4la | 0.5 |
| Area hOc5 [V5/MT] | 0.5 |
| Area FG2 | 0.1 |
| Area FG4 | 0.1 |
| Area FG1 | 0.1 |
| CA1 (Hippocampus) | 0.0 |
| Area hOc1 [V1] | 0.0 |

Probabilities at local maxima -36.0 / -88.0 / +8.0 (Height: 14) -30.0 / -44.0 / -12.0 (Height: 12) -32.0 / -46.0 / -10.0 (Height: 12) -26.0 / -58.0 / -8.0 (Height: 11) -42.0 / -78.0 / -2.0 (Height: 8.8) -30.0 / -74.0 / +0.0 (Height: 7.9) -44.0 / -62.0 / -4.0 (Height: 7.5) -20.0 / -76.0 / -10.0 (Height: 7.4) -22.0 / -78.0 / -8.0 (Height: 7.4)

| | | | | | | | | | |
|---|---|---|---|---|---|---|---|---|---|
| Area hOc4v [V4(v)] | 0.0 | 0.0 | 0.0 | 0.0 | 0.0 | 0.1 | 0.0 | 66.5 | 65.3 |
| Area FG3 | 0.0 | 38.1 | 71.3 | 32.5 | 0.0 | 0.0 | 0.0 | 0.0 | 0.0 |
| Area hOc4lp | 76.4 | 0.0 | 0.0 | 0.0 | 0.0 | 0.0 | 0.0 | 0.0 | 0.0 |
| Area hOc3v [V3v] | 0.0 | 0.0 | 0.0 | 0.0 | 0.0 | 0.0 | 0.0 | 33.1 | 34.2 |
| Area hOc4la | 23.6 | 0.0 | 0.0 | 0.0 | 35.8 | 0.0 | 0.0 | 0.0 | 0.0 |
| Area hOc5 [V5/MT] | 0.0 | 0.0 | 0.0 | 0.0 | 50.3 | 0.0 | 0.0 | 0.0 | 0.0 |
| Area FG2 | 0.0 | 0.0 | 0.0 | 0.0 | 13.9 | 0.0 | 0.0 | 0.0 | 0.0 |
| Area FG4 | 0.0 | 0.0 | 0.0 | 0.0 | 0.0 | 0.0 | 8.2 | 0.0 | 0.0 |
| Area FG1 | 0.0 | 0.0 | 0.0 | 0.0 | 0.0 | 5.8 | 0.0 | 0.5 | 0.5 |
| CA1 (Hippocampus) | 0.0 | 3.1 | 0.0 | 0.0 | 0.0 | 0.0 | 0.0 | 0.0 | 0.0 |
| Area hOc1 [V1] | 0.0 | 0.0 | 0.0 | 0.8 | 0.0 | 0.0 | 0.0 | 0.0 | 0.0 |

Cluster 3 32 vox +18 -62 +52

Assignment based on Maximum Probability Map Percent of Cluster volume in Area Percent of Area activated by Cluster

Area 7A (SPL)33.2 0.6

Area hIP3 (IPS) 10.2 0.3

Area 7P (SPL) 3.9 0.2

Probability exceedance by Area Mean probability at Cluster location (%) Mean probability across PMap (%) Ratio 95% CI 95% CI

Area 7A (SPL)28.0 24.2 1.15 1.07 1.23

Area hIP3 (IPS) 16.5 18.2 0.90 0.79 1.01

Area 7P (SPL) 14.2 22.4 0.63 0.58 0.70

Area 5L (SPL) 5.1 20.1 0.25 0.14 0.38

Area 7PC (SPL) 3.7 17.1 0.22 0.10 0.34

Top probabilites at peak voxels Union across peaks

Area 7A (SPL)0.5

Area hIP3 (IPS) 0.2

Area 7P (SPL) 0.0

Area 5L (SPL) 0.0

Area 7PC (SPL) 0.0

Probabilities at local maxima +18.0 / -62.0 / +52.0 (Height: 6.3) +22.0 / -56.0 / +56.0 (Height: 5.6)

Area 7A (SPL)28.6 36.2

Area hIP3 (IPS) 15.1 3.1

Area 7P (SPL) 4.6 0.0

Area 5L (SPL) 0.0 2.5

Area 7PC (SPL) 0.0 0.6

**t-contrast Faces > Chairs ($p_{FWE}$= .05)**

Image: MAIN_t_Faces-Chairs_FWE05 [u=0.0, k=0]

Cluster 1 64 vox +54 -50 +8

Assignment based on Maximum Probability Map Percent of Cluster volume in Area Percent of Area activated by Cluster

Probability exceedance by Area Mean probability at Cluster location (%) Mean probability across PMap (%) Ratio 95% CI 95% CI

| | | | | | |
|---|---|---|---|---|---|
| Area PGa (IPL) | 4.6 | 20.5 | 0.23 | 0.19 | 0.25 |
| Area PFm (IPL) | 0.6 | 21.7 | 0.03 | 0.00 | 0.06 |

| | | | | | |
|---|---|---|---|---|---|
| Area PGp (IPL) | 1.1 | 25.1 | 0.04 | 0.01 | 0.09 |

| Top probabilites at peak voxels | Union across peaks |
|---|---|
| Area PGa (IPL) | 0.0 |

Probabilities at local maxima +54.0 / -50.0 / +8.0 (Height: 6.5)

| | |
|---|---|
| Area PGa (IPL) | 3.1 |

**t-contrast Distinctive > Typical ($p_{FWE}$= .05)**

Image: MAIN_t_Dist-Typ_FWE05 [u=0.0, k=0]

Cluster 1 293 vox +34 -80 -10

| Assignment based on Maximum Probability Map | Percent of Cluster volume in Area | Percent of Area activated by Cluster |
|---|---|---|
| Area hOc4la | 49.2 | 8.4 |
| Area hOc4v [V4(v)] | 29.6 | 6.4 |
| Area hOc4lp | 10.2 | 2.1 |
| Area hOc3v [V3v] | 5.6 | 1.0 |
| Area FG2 | 1.9 | 0.7 |
| Area FG1 | 1.2 | 0.6 |

| Probability exceedance by Area | Mean probability at Cluster location (%) | Mean probability across PMap (%) | Ratio | 95% CI | 95% CI |
|---|---|---|---|---|---|
| Area hOc4la | 47.4 | 30.4 | 1.56 | 1.52 | 1.61 |
| Area hOc4v [V4(v)] | 29.1 | 25.3 | 1.15 | 1.11 | 1.19 |
| Area hOc4lp | 18.5 | 25.6 | 0.72 | 0.69 | 0.75 |
| Area FG1 | 15.6 | 18.9 | 0.83 | 0.79 | 0.86 |
| Area hOc3v [V3v] | 23.5 | 25.2 | 0.93 | 0.85 | 1.00 |
| Area FG2 | 11.7 | 20.1 | 0.58 | 0.55 | 0.62 |
| Area hOc2 [V2] | 3.0 | 20.6 | 0.15 | 0.10 | 0.19 |
| Area hOc5 [V5/MT] | 1.1 | 15.0 | 0.07 | 0.05 | 0.10 |

| Top probabilites at peak voxels | Union across peaks |
|---|---|
| Area hOc4la | 1.0 |
| Area hOc4v [V4(v)] | 0.9 |
| Area FG1 | 0.5 |

| | |
|---|---|
| Area hOc4lp | 0.3 |
| Area FG2 | 0.1 |
| Area hOc3v [V3v] | 0.0 |
| Area hOc5 [V5/MT] | 0.0 |

Probabilities at local maxima +34.0 / -80.0 / -10.0 (Height: 7.8) +44.0 / -80.0 / +0.0 (Height: 7.6) +36.0 / -78.0 / -12.0 (Height: 7.6) +42.0 / -78.0 / -2.0 (Height: 7.6)

| | | | | |
|---|---|---|---|---|
| Area hOc4la | 0.0 | 82.2 | 0.0 | 74.6 |
| Area hOc4v [V4(v)] | 65.1 | 0.0 | 61.8 | 0.0 |
| Area FG1 | 22.7 | 0.0 | 31.5 | 0.0 |
| Area hOc4lp | 3.3 | 17.7 | 0.7 | 10.7 |
| Area FG2 | 0.0 | 0.0 | 6.0 | 0.0 |
| Area hOc3v [V3v] | 4.6 | 0.0 | 0.0 | 0.0 |
| Area hOc5 [V5/MT] | 0.0 | 0.0 | 0.0 | 0.0 |

Cluster 2 159 vox -34 -86 -6

Assignment based on Maximum Probability Map Percent of Cluster volume in Area Percent of Area activated by Cluster

| | | |
|---|---|---|
| Area hOc4v [V4(v)] | 31.4 | 3.7 |
| Area hOc4la | 28.7 | 2.7 |
| Area FG2 | 11.0 | 2.2 |
| Area hOc3v [V3v] | 10.7 | 1.0 |
| Area hOc4lp | 8.8 | 1.0 |

| | | |
|---|---|---|
| Area FG1 | 0.2 | 0.0 |
| Area hOc5 [V5/MT] | 0.1 | 0.1 |

| Probability exceedance by Area | Mean probability at Cluster location (%) | Mean probability across PMap (%) | Ratio | 95% CI | 95% CI |
|---|---|---|---|---|---|
| Area hOc4v [V4(v)] | 33.1 | 25.3 | 1.31 | 1.25 | 1.37 |
| Area hOc4la | 28.7 | 30.4 | 0.94 | 0.89 | 0.99 |
| Area hOc3v [V3v] | 22.8 | 25.2 | 0.90 | 0.84 | 0.95 |
| Area FG2 | 19.9 | 20.1 | 0.99 | 0.93 | 1.05 |
| Area hOc4lp | 19.7 | 25.6 | 0.77 | 0.70 | 0.82 |
| Area FG1 | 7.9 | 18.9 | 0.42 | 0.39 | 0.45 |
| Area hOc5 [V5/MT] | 6.7 | 15.0 | 0.45 | 0.38 | 0.52 |
| Area hOc2 [V2] | 3.0 | 20.6 | 0.15 | 0.13 | 0.17 |
| Area FG4 | 0.7 | 24.1 | 0.03 | 0.01 | 0.05 |

| Top probabilites at peak voxels | Union across peaks |
|---|---|
| Area hOc4v [V4(v)] | 0.9 |
| Area hOc4la | 0.9 |
| Area FG2 | 0.5 |
| Area hOc3v [V3v] | 0.4 |
| Area hOc4lp | 0.4 |
| Area FG1 | 0.2 |
| Area hOc5 [V5/MT] | 0.1 |
| Area hOc2 [V2] | 0.0 |

| Probabilities at local maxima | -34.0 / -86.0 / -6.0 (Height: 7.1) | -24.0 / -86.0 / -12.0 (Height: 6.7) | -28.0 / -84.0 / -12.0 (Height: 6.7) | -40.0 / -72.0 / -8.0 (Height: 6.2) | -40.0 / -78.0 / -6.0 (Height: 6.1) | -44.0 / -80.0 / -8.0 (Height: 6.1) |
|---|---|---|---|---|---|---|
| Area hOc4v [V4(v)] | 22.6 | 62.5 | 74.3 | 0.0 | 0.2 | 0.0 |
| Area hOc4la | 32.8 | 0.0 | 0.0 | 9.9 | 18.4 | 72.4 |
| Area FG2 | 0.0 | 0.0 | 0.0 | 19.3 | 27.3 | 14.6 |

| | | | | | | |
|---|---|---|---|---|---|---|
| Area hOc3v [V3v] | 3.3 | 36.4 | 7.0 | 0.0 | 0.0 | 0.0 |
| Area hOc4lp | 37.6 | 0.0 | 0.0 | 0.0 | 0.0 | 0.9 |
| Area FG1 | 2.5 | 0.0 | 2.4 | 9.2 | 6.9 | 0.0 |
| Area hOc5 [V5/MT] | 0.0 | 0.0 | 0.0 | 0.0 | 2.7 | 5.3 |
| Area hOc2 [V2] | 0.0 | 1.0 | 0.0 | 0.0 | 0.0 | 0.0 |

Cluster 3 3 vox +40 -60 -12

| Assignment based on Maximum Probability Map | Percent of Cluster volume in Area | Percent of Area activated by Cluster |
|---|---|---|
| Area FG2 | 62.5 | 0.2 |
| Area FG1 | 33.3 | 0.2 |

| Probability exceedance by Area | Mean probability at Cluster location (%) | Mean probability across PMap (%) | Ratio | 95% CI | 95% CI |
|---|---|---|---|---|---|
| Area FG2 | 30.0 | 20.1 | 1.49 | 1.33 | 1.65 |
| Area FG1 | 25.3 | 18.9 | 1.34 | 1.09 | 1.56 |
| Area FG4 | 16.6 | 24.1 | 0.69 | 0.60 | 0.77 |
| Area hOc4la | 2.0 | 30.4 | 0.07 | 0.03 | 0.09 |
| Area hOc4v [V4(v)] | 0.1 | 25.3 | 0.01 | 0.01 | 0.01 |

| Top probabilites at peak voxels | | Union across peaks |
|---|---|---|
| Area FG1 | 0.3 | |
| Area FG4 | 0.2 | |
| Area FG2 | 0.2 | |

Probabilities at local maxima +40.0 / -60.0 / -12.0 (Height: 5.7)

| | |
|---|---|
| Area FG1 | 27.6 |
| Area FG4 | 19.0 |
| Area FG2 | 16.9 |

Cluster 4 1 vox +46 -58 -12

Assignment based on Maximum Probability Map Percent of Cluster volume in Area Percent of Area activated by Cluster

Probability exceedance by Area Mean probability at Cluster location (%) Mean probability across PMap (%) Ratio 95% CI 95% CI

| | | | | | |
|---|---|---|---|---|---|
| Area FG2 | 18.5 | 20.1 | 0.92 | 0.82 | 1.03 |
| Area FG4 | 12.3 | 24.1 | 0.51 | 0.44 | 0.57 |
| Area hOc4la | 2.6 | 30.4 | 0.08 | 0.05 | 0.11 |

Top probabilites at peak voxels Union across peaks

| | |
|---|---|
| Area FG4 | 0.1 |
| Area FG2 | 0.1 |

Probabilities at local maxima +46.0 / -58.0 / -12.0 (Height: 5.7)

| | |
|---|---|
| Area FG4 | 13.1 |
| Area FG2 | 13.1 |

---

**t-contrast Typical > Distinctive ($p_{FWE}$= .05)**

---

Image: Opposite_t_Typ-Dist_FWE05 [u=0.0, k=0]

Cluster 1 8 vox -4 -64 +24

Assignment based on Maximum Probability Map Percent of Cluster volume in Area Percent of Area activated by Cluster

Probability exceedance by Area Mean probability at Cluster location (%) Mean probability across PMap (%) Ratio 95% CI 95% CI

Area 7A (SPL)1.8 24.2 0.07 0.04 0.12

Top probabilites at peak voxels Union across peaks

Area 7A (SPL)0.0

Probabilities at local maxima -4.0 / -64.0 / +24.0 (Height: 6.1)

Area 7A (SPL)0.0

Cluster 2 4 vox -8 -46 +36

Assignment based on Maximum Probability Map Percent of Cluster volume in Area Percent of Area activated by Cluster

Probability exceedance by Area Mean probability at Cluster location (%) Mean probability across PMap (%) Ratio 95% CI 95% CI

Top probabilites at peak voxels Union across peaks

Probabilities at local maxima

Reference List

**t-contrast High Pred Faces AND High Pred Chairs > Uninformative ($p_{unc.}$= .0001)**

Image: MAIN_t_highExp-uninf_unc0001 [u=0.0, k=0]

Cluster 1 3 vox +10 +4 +26

Assignment based on Maximum Probability Map Percent of Cluster volume in Area Percent of Area activated by Cluster

Probability exceedance by Area	Mean probability at Cluster location (%)	Mean probability across PMap (%)	Ratio	95% CI	95% CI

Area 33	1.9	17.5	0.11	0.06	0.16

Top probabilites at peak voxels	Union across peaks

Probabilities at local maxima

Cluster 2	1 vox	-10	-2	+30

Assignment based on Maximum Probability Map	Percent of Cluster volume in Area	Percent of Area activated by Cluster

Area 33	100.0	0.3

Probability exceedance by Area	Mean probability at Cluster location (%)	Mean probability across PMap (%)	Ratio	95% CI	95% CI

Area 33	55.2	17.5	3.15	2.80	3.51

Top probabilites at peak voxels	Union across peaks

Area 33	0.5

Probabilities at local maxima -10.0 / -2.0 / +30.0 (Height: 3.6)

Area 33	46.6

**F-contrast Predictability ($p_{unc.}$= .0001)**

Image: MAIN_f_Pred_unc0001 [u=0.0, k=0]

Cluster 1 14 vox +10 +4 +26

Assignment based on Maximum Probability Map Percent of Cluster volume in Area Percent of Area activated by Cluster

Probability exceedance by Area Mean probability at Cluster location (%) Mean probability across PMap (%) Ratio 95% CI 95% CI

| | | | | | |
|---|---|---|---|---|---|
| Area 33 | 5.3 | 17.5 | 0.30 | 0.21 | 0.40 |

Top probabilites at peak voxels Union across peaks

Probabilities at local maxima

Cluster 2 12 vox -10 -2 +30

Assignment based on Maximum Probability Map Percent of Cluster volume in Area Percent of Area activated by Cluster

Area 33 50.0 1.6

Probability exceedance by Area Mean probability at Cluster location (%) Mean probability across PMap (%) Ratio 95% CI 95% CI

Area 33 38.5 17.5 2.20 1.96 2.45

Top probabilites at peak voxels Union across peaks

Area 33 0.5

Probabilities at local maxima -10.0 / -2.0 / +30.0 (Height: 18)

Area 33 46.6

Cluster 3 1 vox -40 -28 -2

Assignment based on Maximum Probability Map Percent of Cluster volume in Area Percent of Area activated by Cluster

Probability exceedance by Area Mean probability at Cluster location (%) Mean probability across PMap (%) Ratio 95% CI 95% CI

| | | | | | |
|---|---|---|---|---|---|
| Area Id1 | 7.9 | 15.9 | 0.50 | 0.20 | 0.79 |
| Area Ig1 | 0.7 | 15.4 | 0.05 | 0.02 | 0.08 |

Top probabilites at peak voxels Union across peaks

| | |
|---|---|
| Area Id1 | 0.2 |
| Area Ig1 | 0.0 |

Probabilities at local maxima -40.0 / -28.0 / -2.0 (Height: 13)

| | |
|---|---|
| Area Id1 | 19.2 |
| Area Ig1 | 0.3 |

**F-contrast Predictability x Typicality Interaction ($p_{unc.}$= .0001)**

No significant results
