## Supplementary materials for "Investigating the neural effects of typicality and predictability for face and object stimuli"

Table of contents:

*Table S1.* Summary of post-experimental survey	**2**

*Table S2.* ROI locations in individual participants	**4**

***Table S3.*** Contrast weights for the whole-brain analysis	**6**

***Figure S1.*** ANOVA results in individual ROIs	**7**

***Figure S2.*** Whole-brain typicality effects for faces and chairs separately	**8**

Table S1. **Summary of post-experimental survey**

Table S1: Synopsis of the post-experimental survey completed by participants Each participant was first asked about whether they experienced any form of discomfort during the procedure, and eventually which ones. Then they were asked to indicate to which degree they found themselves paying attention to cue-category contingencies during the scanning session. Their verbal reports (e.g., “never”, not really”, “at the beginning only”, “sometimes”, etc.) were then transferred to a scale ranging from 0 (= “not at all”) to 3 (= “very frequently/always”). No participant reported a score of 3. Participants were then asked whether the faces and the chairs shown during the tasks look ordinary to them or whether they had something distinctive. Their responses were coded as “yes” if they made observations about differential typicality within that category (e.g., “some of the chairs were quite special”), and with “no” if all items of that category appeared of equal typicality to them (e.g., “all faces looked pretty normal to me, none stood out”). As shown in the total proportions of “yes” responses, typicality manipulations were detected much more often in chairs than faces.

| **SubjectID** | **Discomforts during scanning?** | **Which discomforts?** | **Attention to cue-stimulus contingencies** | **Typicality manipulation in objects?** | **Typicality manipulation in faces?** |
| --- | --- | --- | --- | --- | --- |
| NiKr92 | Yes | itching, remaining still | 1 | Yes | No |
| JoMe03 | Yes | earplugs, head adjustment | 1 | No | No |
| ZiHe98 | Yes | glasses | 1 | Yes | No |
| JoBi01 | No | none | 0 | No | No |
| KeSi01 | No | none | 0 | No | Yes |
| RiJa02 | Yes | head pain | 1 | No | No |
| FeBa02 | No | none | 1 | Yes | No |
| AmKa01 | Yes | staying still, boredom | 2 | Yes | No |
| MoZe01 | Yes | hard to stay focussed | 1 | No | No |
| UlKr94 | Yes | numb fingers | 0 | No | No |
| MiWe97 | Yes | tired | 2 | No | No |
| KaSo01 | Yes | tired, boredom | 1 | No | No |
| PiEl00 | Yes | eye strain right | 2 | No | No |
| PaPo97 | Yes | head pain | 0 | Yes | No |
| KlBeHe01 | No | none | 0 | No | No |
| RoTe91 | No | none | 2 | Yes | No |
| MeBo00 | Yes | tired | 2 | Yes | No |
| LeFi01 | No | none | 0 | Yes | No |
| SiLa01 | No | none | 2 | Yes | Yes |
| PaAlHa96 | Yes | foot pain, tired | 0 | Yes | No |
| ViKl03 | No | none | 2 | No | No |
| **SubjectID** | **Discomforts during scanning?** | **Which discomforts?** | **Attention to cue-stimulus contingencies** | **Typicality manipulation in objects?** | **Typicality manipulation in faces?** |
| LiHo94 | Yes | vibration, tired | 0 | Yes | Yes |
| LaSp91 | Yes | head pain | 0 | Yes | No |
| SeHu99 | Yes | tired | 2 | Yes | No |
| ViIv95 | Yes | tired | 0 | Yes | No |
| RiSi95 | No | none | 1 | Yes | No |
| AmAi98 | Yes | head pain, eye strain | 0 | Yes | No |
| ChRz87 | Yes | head pain | 0 | No | No |
| AnFi01 | No | none | 0 | No | No |
| IlCh95 | Yes | tired, cold | 0 | Yes | No |
| NaSc91 | Yes | earplugs uncomfortable | 0 | Yes | No |
| ReRu02 | No | none | 2 | Yes | No |
| YaEi97 | No | none | 0 | No | No |
| ViPa00 | Yes | boredom, head pain | 1 | Yes | Yes |
| JoKn87 | Yes | cold, tingling, tired | 0 | No | Yes |
| **Summary** | **22/35 “yes”** | **-** | **M = 0,77 (SD = 0,84)** | **20/35 “yes”** | **5/35 “yes”** |

Table S2. **ROI locations in individual participants**

Table S2: MNI coordinates of activation peaks in individual participants during functional localizer block. Right and left fusiform faces areas (FFAs) were localized using the contrast Faces > Objects (or Faces > Noise when the first did not reveal any peak in the approximate area). Right and left lateral occipital complexes (LOCs) were localized using the contrast Objects > Noise. Stars (*) indicate that the coordinate was retrieved when changing the threshold from $p_{FWE}<.05$ to $p_{uncorrected}<.001$. In a few cases no peaks were found for a particular area. When comparing this table to Fig. S1 and Fig. 3, note that even though we were able to locate these ROIs in nearly all participants, at times the hemodynamic function extracted at the respective location was not of sufficient quality, thus data were excluded from the analysis reported in the main text.

| **SubjectID** | **Right FFA** | **Left FFA** | **Right LOC** | **Left LOC** |
| --- | --- | --- | --- | --- |
| NiKr92 | 42, -58, -14 | -44, -52, -22* | 28,-86, -16 | -36, -80, 8 |
| JoMe03 | Not found | -38, -48, -18* | 40, -78, -2 | -50, -72, 0 |
| ZiHe98 | 42, -64, -14* | -44, -62, -16* | 40, -86, 2 | -36, -74, -2 |
| JoBi01 | 48, -58, -29* | -46, -50, -20* | 44, -78, 2 | -38, -90, 8 |
| KeSi01 | 36, -46, -20 | -32, -56, -18 | 38, -72, 0 | -46, -80, 14 |
| RiJa02 | 34, -50, -20* | Not found | 42, -84, -8* | -52, -70, -12 |
| FeBa02 | 40, -60, -14 | -42, -66, -16 | 38, -78, -2 | -48, -82, 0 |
| AmKa01 | 36, -60, -16 | -38, 56, -18 | 38, -92, 4 | -42, -78, 8 |
| MoZe01 | 42, -58, -20 | -44, -62, -18* | 34, -78, 4 | -42, -76, -10 |
| UlKr94 | 42, -42, -18* | -42, -48, -20 | 42, -86, 2 | -36, -92, -2 |
| MiWe97 | 34, -64, -14* | -38, -48, -22* | 46, -72, -2 | -44, -84, -6 |
| KaSo01 | 44, -46, -16 | -44, -46, -18 | 48, -74, 4 | -40, -74, 0 |
| PiEl00 | 40, -54, -20 | -40, -60, -22 | 40, -76, 2 | -50, -76, -8 |
| PaPo97 | 38, -56, -16* | -32, -56, -16 | 38, -82, -8 | -36, -82, -2 |
| KlBeHe01 | 44, -46, -20* | -40, -52, -20* | 44, -66, -6 | -52, -70, -2 |
| RoTe91 | 42, -64, -14 | -42, -52, -22 | 38, -82, -4 | -46, -76, -6 |
| MeBo00 | 36, -48, -20 | -38, -46, -18 | 32, -94, 8 | -38, -90, 4 |
| LeFi01 | 38, -48, -22* | -38, -56, -20 | 40, -78, -4 | -48, -72, 0 |
| SiLa01 | 40, -52, -18* | -38, -52, -20* | 46, -80, -4 | -42, 72, -12 |
| PaAlHa96 | 46, -64, -20 | -38, -46, -22 | 50, -78, -4 | -50, -70, -4 |
| ViKl03 | 46, -52, -16 | -38 -56, -16* | 52, -70, -8 | -34, -84, 0 |
| LiHo94 | 36, -48, -20 | -34, -48, -24 | 50, -70, 2 | -48, -82, -6 |
| LaSp91 | 36, -40, -24* | -38, -42, -16* | 40, -86, 2 | -48, -80, -10 |
| SeHu99 | 42, -44, -14 | -36, -66, -4 | 46, -78, -14 | -48, -78, 0 |
| ViIv95 | 54, -66, -14 | Not found | 50, -80, 6 | -48, -82, -4 |
| **SubjectID** | **Right FFA** | **Left FFA** | **Right LOC** | **Left LOC** |
| RiSi95 | 44, -58, -22 | -38, -40, -26 | 46, -80, 6 | -4, -80, 2 |
| AmAi98 | 48, -54, -24* | -40, -48, -20* | 44, -84, -14 | -42, -86, -8 |
| ChRz87 | 48, -62, -18 | -42, -58, -16* | 40, -70, -6* | -38, -72, -10* |
| AnFi01 | 38, -56, -16 | -36, -62, -16 | 48, -86, 2 | -46, -76, -2 |
| IlCh95 | 36, -48, -14 | -38, -54, -18 | 46, -80, -4 | -48, -78, -6 |
| NaSc91 | 38, -46, -24* | -38, -44, -16* | 44, -80, -4 | -40, -72, -12 |
| ReRu02 | 36, -52, -16 | -32, -58, -20* | 42, -86, 4 | -46, -78, 0 |
| YaEi97 | 38, -46, -18 | -38, -48, -24 | 32, -88, 6 | -46, -80, -4 |
| ViPa00 | 40, -42, -26 | -40, -44, -26 | 40, -82, -2 | -42, -76, -8 |
| JoKn87 | 38, -5, -18* | -38, -52, -18 | 42, -78, -1 | -44, -78, -4 |
| **Average(SD)** | **x = 40,6(4,7)**  **y = -52,9(7,7)**  **z = -18,6(3,9)** | **x = -38,9(3,5)**  **y = -52,4(6,9)**  **z = -18,9(4,0)** | **x = 40,1(4,6)**  **y = -76,1(8,2)**  **z = -3,6(6,4)** | **x = -43,5(5,1)**  **y = -78,3(6,6)**  **z = -3,5(5,7)** |

Table S3. Contrast weights for the whole-brain analysis

*Table S3: Conditions are abbreviated as follows: H = high predictability cue (75% contingencies); M = medium/uninformative cue (50% contingencies); L = low predictability cue (25 % contingencies); F = face stimulus; C = chair stimulus; T = typical stimulus; D = distinctive stimulus. Note that weights are set to 0 for target trials and for the six nuisance regressors.*

| **Contrast weights for the whole brain analysis** **Conditions** H_F_T \| H_F_D \| H_C_T \| H_C_D \| M_F_T \| M_F_D \| M_C_T \| M_C_D \| L_F_T \| L_F_D \| L_C_T \| L_C_D \| Target_trials and six nuisance regressors | |
| --- | --- |
| Main effects of category | t-contrast: Faces > Chairs [1 1 -1 -1 1 1 -1 -1 1 1 -1 -1 0 0 0 0 0 0 0]  t-contrast: Chairs > Faces [-1 -1 1 1 -1 -1 1 1 -1 -1 1 1 0 0 0 0 0 0 0] |
| Main effects of typicality | t-contrast: Distinctive > Typical [-1 1 -1 1 -1 1 -1 1 -1 1 -1 1 0 0 0 0 0 0 0]  t-contrast: Typical > Distinctive [1 -1 1 -1 1 -1 1 -1 1 -1 1 -1 0 0 0 0 0 0 0] |
| Main effects of predictability | Directional effect (t-contrast: High Face Predictability and High Chair Predictability > Uninformative Cue)  [1 1 1 1 -2 -2 -2 -2 1 1 1 1 0 0 0 0 0 0 0]  Non-directional effect (f-contrast: [1 1 1 1 -1 -1 -1 -1 0 0 0 0 0 0 0 0 0 0 0  0 0 0 0 -1 -1 -1 -1 1 1 1 1 0 0 0 0 0 0 0]) |
| Interaction between typicality and predictability | f-contrast: [1 -1 1 -1 -1 1 -1 1 0 0 0 0 0 0 0 0 0 0 0  0 0 0 0 1 -1 1 -1 -1 1 -1 1 0 0 0 0 0 0 0] |

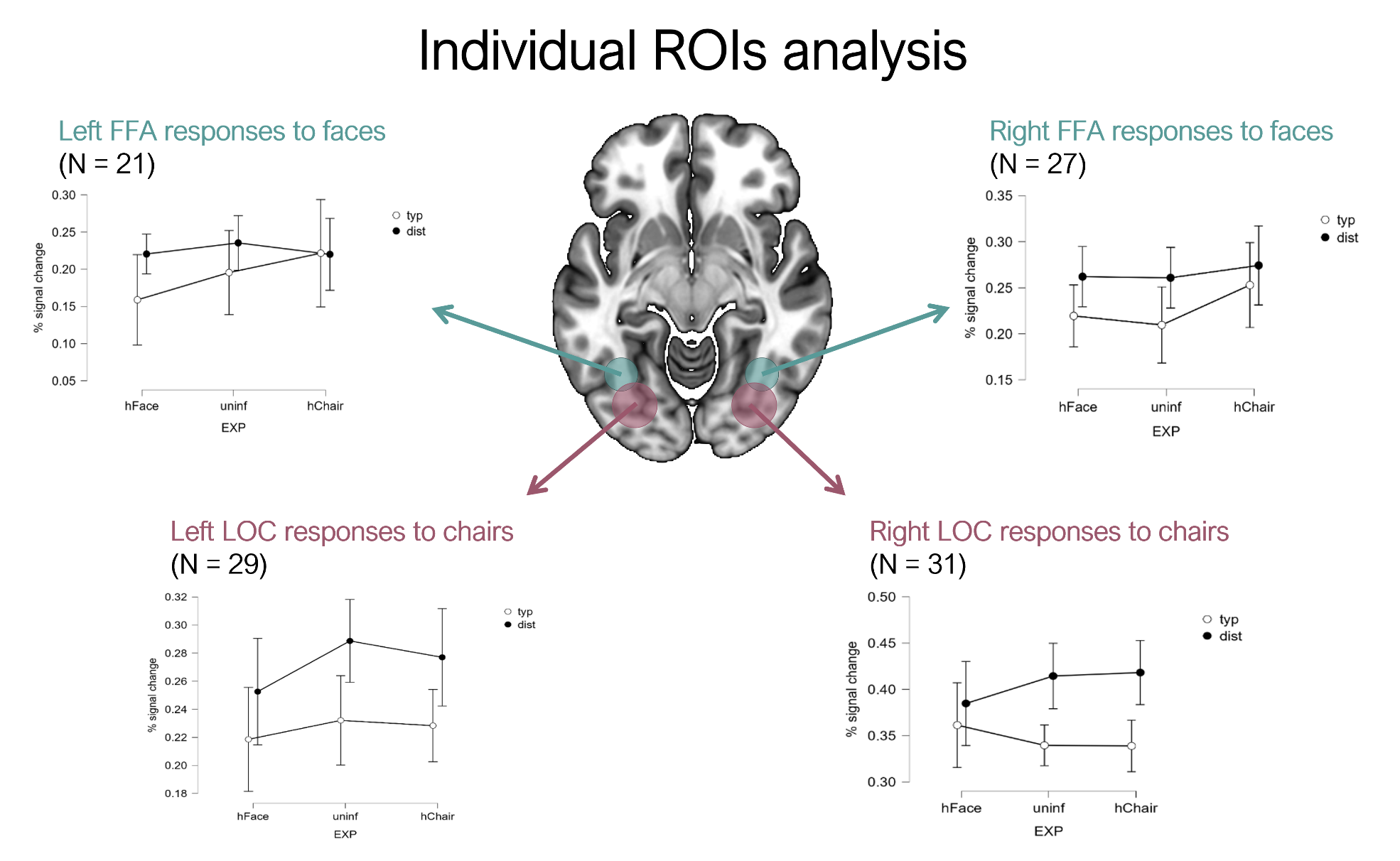
Figure S1. **ANOVA results in individual ROIs**

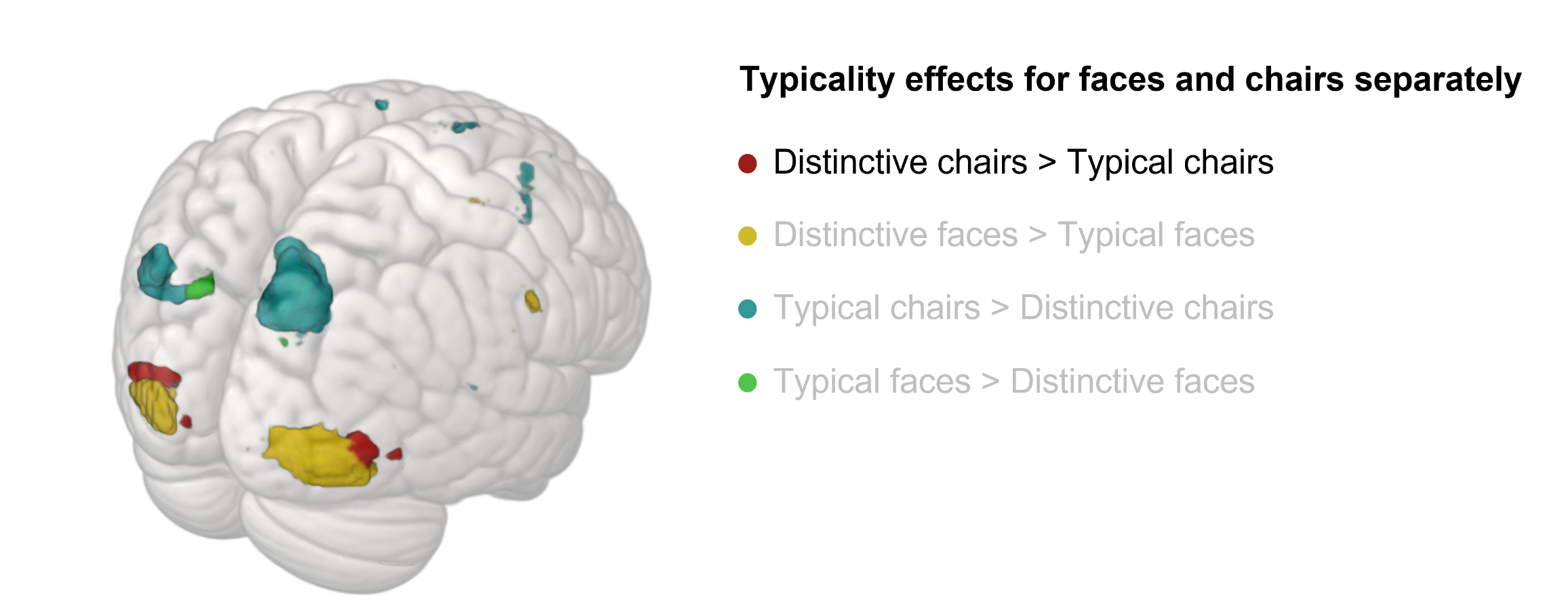
**Figure S2: Whole-brain typicality effects for faces and chairs separately**

Figure S2: whole-brain effects of typicality in faces and chairs separately (directional t-contrasts). Results for the contrast distinctive chairs > typical chairs are thresholded at an alpha level of .05 and a family-wise error correction for multiple comparison (FWE, p < .05). The other contrasts are thresholded at an alpha level of .001 and no correction (uncorrected, p < .001). The ink transparency in the legend visually conveys the different thresholding conservativeness of the results. This figure shows that the effects of distinctiveness (distinctive > typical) tend to occur in overlapping regions for both stimulus types (note that, with an uncorrected threshold, the clusters related to chair distinctiveness overlap with those related to face distinctiveness). Conversely, typicality effects seem to be differently distributed and to be more prominent for chairs.
